# Age- and episodic memory-related differences in task-based functional connectivity in women and men

**DOI:** 10.1101/2021.07.27.453878

**Authors:** Sivaniya Subramaniapillai, Sricharana Rajagopal, Elizabeth Ankudowich, Stamatoula Pasvanis, Bratislav Misic, M.Natasha Rajah

## Abstract

Aging is associated with episodic memory decline and changes in functional brain connectivity. Understanding whether and how biological sex influences age- and memory performance-related functional connectivity has important theoretical and clinical implications for our understanding of brain and cognitive aging. Yet, little is known about the effect of sex on neurocognitive aging. Here, we scanned 161 healthy adults between 19-76 yrs of age in an event-related functional magnetic resonance imaging (fMRI) study of face-location spatial context memory. Adults were scanned while performing easy and difficult versions of the task at both encoding and retrieval. We used multivariate whole-brain partial least squares (PLS) connectivity to test the hypothesis that there are sex differences in age- and episodic memory performance-related functional connectivity. We examined how individual differences in age and retrieval accuracy correlated with task-related connectivity. We then repeated this analysis after disaggregating the data by self-reported sex. We found that increased encoding and retrieval-related connectivity within the dorsal attention network (DAN), and between DAN and frontoparietal network (FPN) and visual networks, was positively correlated to retrieval accuracy and negatively correlated with age in both sexes. We also observed sex differences in age- and performance-related functional connectivity: i) greater between-network integration was apparent at both levels of task difficulty in women only, and ii) increased DAN-DMN connectivity with age was observed in men and was correlated with poorer memory performance. Therefore, the neural correlates of age-related episodic memory decline differ in women and men and has important theoretical and clinical implications for the cognitive neuroscience of memory, aging and dementia prevention.

## Introduction

Healthy aging is associated with episodic memory decline, a reduced ability to encode, store and retrieve past experiences in rich spatio-temporal contextual detail (Grady & Craik, 2000; Tulving, 1972). Age-associated episodic memory decline impairs older adults’ quality of life and can be an early sign of sporadic Alzheimer’s disease (AD) (Mol et al., 2007; Mol, van Boxtel, Willems, & Jolles, 2006). Given that the proportion of older adults is increasing worldwide, and age is the strongest predictor of AD, there is an urgent need to understand how normative aging influences memory and related brain function.

To this aim, there is a large body of research that has investigated how normative aging affects episodic memory and related brain activity using task functional magnetic resonance imaging (fMRI) (Grady, 2008; Maillet & Rajah, 2014; Naveh-Benjamin et al., 2003; Nyberg et al., 2012; Rajah & McIntosh, 2005; Spaniol et al., 2009; Sperling, 2007). This research has shown that age-related reductions in episodic memory, as measured by associative memory tasks (e.g. spatial context memory tasks), are present at midlife and increase with advanced age (Ankudowich, Pasvanis, & Rajah, 2016; Cansino, 2009; Kwon et al., 2016), and that these behavioral reductions are associated with altered activation in occipito-temporal, prefrontal cortex (PFC), inferior parietal cortex (IPC) and medial temporal lobe (MTL) with age (Ankudowich et al., 2016; 2017; 2019). Furthermore, with the growing consensus that human cognition and behavior depends on the dynamic interactions of large-scale neural networks (Friston, 1994; McIntosh, 2000; Mesulam, 1990; Sporns & Betzel, 2016; Strother et al., 1995), several cognitive neuroscience studies of aging have focused on how age differences in inter-regional or inter-network correlations in brain activity (functional connectivity) during resting state fMRI (rsfMRI) relate to cognitive task performance assessed outside of the scanner (Biswal et al., 1995; Power et al., 2011; Yeo et al., 2011; Uddin, Yeo, & Spreng, 2019).

Studies of rsfMRI connectivity have found that age-related decreases in cognitive task performance were associated with reduced anticorrelation between the dorsal attention network (DAN) and default mode network (DMN), possibly as a consequence of disrupted frontoparietal network (FPN) engagement (Amer, Campbell, & Hasher, 2016; Avelar-Pereira et al., 2017; Dixon et al., 2017; Esposito et al., 2018; Fox et al., 2005; Grady et al., 2016; Prakash et al., 2012; Sala-Llonch et al., 2012; Spreng et al., 2016). More generally, aging has also been correlated with increased connectivity between networks (i.e., network integration) and decreased connectivity within networks (i.e., network segregation) (Chan et al., 2014; Damoiseaux, 2017). However, only a few rsfMRI studies have directly explored whether age-related differences in connectivity correlated with pre-/post-scan performance on *episodic memory* tasks (Edde et al., 2020; Fjell et al., 2015; Grady et al., 2016; King et al., 2018; Kukolja et al., 2016; Nordin et al., 2021; Nyberg, 2017; Wang et al., 2010; Zhang, et al., 2020). Most of these studies focused on specific *a priori* defined networks of interest (but see Fjell et al., 2015). Therefore, there remains a paucity of knowledge about how age-related differences in *whole-brain* functional connectivity contribute to decreases in episodic memory with age. Moreover, most of what we know about the correlation between age-related differences in functional connectivity and episodic memory is based on rsfMRI paradigms. Thus, while resting-state research has provided a greater understanding of functional architecture, solely relying on resting state scans as an indirect proxy for cognitive processes is not sufficient to understand brain-cognitive processes (see reviews by Campbell & Schacter, 2016; Finn, 2021).

To our knowledge no prior work has specifically investigated how age and performance correlates with whole brain, task-based functional connectivity during episodic encoding and retrieval, across the adult lifespan. One recent study investigated age-related differences in whole-brain connectivity during encoding of an associative memory task across the adult lifespan (Capogna et al., 2022). Using a whole-brain psychophysiological interaction analysis to investigate direct brain-cognitive processes, the authors found that in older age, greater connectivity between medial temporal and posterior parietal regions during encoding was associated with better performance, while increased connectivity between frontal, parietal, and visual regions was associated with worse performance. The functional connectivity patterns associated with successful memory performance in older adults are associated with cognitive processes that involve integrative and multisensory strategies and mental imagery. However, this study controlled for sex in their analyses hindering any further interpretations of how these findings may separately relate to women and men.

Indeed, most fMRI connectivity studies of aging have assumed that age-related differences in functional connectivity were the same in women and men, since data were not disaggregated by sex and/or gender at analysis. However, depending on the task stimuli and design, studies have repeatedly demonstrated behavioral sex differences on episodic memory performance. Women typically perform better than men on episodic memory tasks of verbal stimuli (Gur & Gur, 2002; Herlitz, Nilsson, & Bäckman, 1997; Ragland, Coleman, Gur, Glahn, & Gur, 2000), whereas men tend perform better than women on visuospatial memory tasks (De Frias, Nilsson, & Herlitz, 2006; Weiss, Kemmler, Deisenhammer, Fleischhacker, & Delazer, 2003). However, these sex differences have small to medium effect sizes and are stable across the adult lifespan (Asperholm, Van Leuven, & Herlitz, 2020; De Frias et al., 2006; Jack et al., 2015; Voyer, Postma, Brake, & Imperato-McGinley, 2007). This may account for the few studies investigating sex differences in age effects on memory and associated brain activity and connectivity. However, even if there are no significant sex and/or sex-by-age interactions in behavioral outcomes, sex differences in the underlying neural system supporting episodic memory across the adult lifespan may still exist (Becker & Koob, 2016; McCarthy, Arnold, Ball, Blaustein, & de Vries, 2012). Consistent with the view that there may be sexual divergence in the brain systems supporting episodic memory function in older women and men, recent studies have found that age-related memory decline was correlated with different patterns of activations in women compared to men (Rabipour et al., 2021; Subramaniapillai et al., 2019). Yet, it remains unclear if there are sex differences in how age and memory performance correlate with task-based functional connectivity during episodic memory encoding and retrieval. This information is important to know because historically it has been assumed that the neural basis of age-associated memory decline is the same in both sexes, but this may not be the case (Ferretti et al., 2018; Nebel et al., 2018; Rahman et al., 2020; Snyder et al., 2016; Subramaniapillai et al., 2021). Investigating sex and gender differences in functional brain connectivity in a normative adult lifespan sample can help determine if there are sex and/or gender-specific markers of memory decline in the aging brain. Such knowledge informs us if the underlying neurocognitive mechanisms linked to age-related episodic memory decline is the same in women and men, and if interventions aimed at supporting memory into late life should be the same for women and men.

Here, we present whole brain functional connectivity results from an episodic memory task fMRI study of 161 healthy adults aged 19 -76 yrs of age who were scanned while performing both encoding and retrieval phases of a face-location spatial context memory paradigm. We parcellated task fMRI data into canonical brain networks defined by Power et al. (2011) and used whole-brain behavior partial least squares (B-PLS) connectivity analysis to examine the orthogonalized contributions of age and memory performance on task-based functional connectivity. We then repeated this analysis after disaggregating the data by self-reported sex to investigate whether both sexes exhibited similar age- and performance-related patterns of connectivity. We hypothesized that age would be correlated with decreased connectivity between DAN – FPN and increased connectivity between DAN – DMN, and memory performance would exhibit the opposite patterns of network associations (Amer et al., 2016; Avelar-Pereira et al., 2017; Dixon et al., 2017; Esposito et al., 2018; Fox et al., 2005; Grady et al., 2016; Prakash et al., 2012; Sala-Llonch et al., 2012; Spreng et al., 2016; Turner & Spreng, 2012). Based on prior activation analyses of sex differences in the effect of age and memory accuracy on task-related brain activity across the adult lifespan (Subramaniapillai et al., 2019), we also hypothesized that both sexes will exhibit similar patterns of performance-related functional connectivity at encoding, but not retrieval. We also hypothesized that there would be sex differences in age-related functional connectivity at both encoding and retrieval.

## Methods

### Participants

Volunteer research participants were recruited from the Montreal and surrounding area using online and print advertisements and community outreach. Research volunteers were told they would first be asked to participate in a behavioral and neuropsychological testing session (Visit 1), and if they met our inclusion criteria, they would be invited back for an fMRI session (Visit 2). Two hundred and seventy-five participants (102 self-identified as men, 173 self-identified as women) were tested in Visit 1. Of these, 49 were excluded for not meeting our neuropsychological inclusion criteria (listed below), 26 were excluded for having medical/psychiatric exclusionary criteria (listed below), and 15 participants could not be reached for scheduling a Visit 2. Therefore, 185 participants were invited back for Visit 2 and participated in the fMRI portion of this study. Of these participants, we identified incidental findings in 9 participants, 5 participants fMRI data did not meet our quality control criteria (listed below), and 10 participants did not perform the fMRI task as instructed, resulting in a sample of 161 participants (49 men, 112 women) who reported no history of neurological or psychological illness, or serious cardiovascular disease. All participants were right-handed, as confirmed by the Edinburgh Inventory for Handedness. Of the 53 middle-aged women, we had self-reported menopause status for 41 women, 18 of these self-reported having irregular periods, symptoms of the menopausal transition, and/or had undergone hormone replacement therapy (HRT). Two older adult women had also undergone HRT. Thus, we excluded these 20 women from further analyses since menopause transition and HRT influences memory-related brain activity (Henderson, 2010; Li, Cui, & Shen, 2014; Rentz et al., 2017; Yonker et al., 2006). Our final cohort consisted of 141 participants (49 men, 92 women; 65% women) between the ages of 19 -76 yrs (mean age = 47.11, SE = 1.41, mean education = 15.73 yrs, SE = 0.18). Of the 35 middle-aged women, we had a self-reported pre-menopausal status for 23 women, with unknown status for 12 women. As we did not have hormonal data to verify self-reported menopausal status, we focus here on age and sex effects and note in our Caveats the need to consider reproductive age and health in future studies examining sex differences in brain aging.

### Behavioral Methods

#### Visit 1: Behavioral and Neuropsychological Session

During an initial session, participants provided informed consent and then were administered a medical screening questionnaire to assess neurological, psychological, and physical health. Medical health exclusion criteria for this study included having a current diagnosis of diabetes, untreated cataracts and glaucoma, and a current diagnosis of high cholesterol levels and/or high blood pressure left untreated in past 6 months. In addition, participants were excluded if they had a history of a major psychiatric illness, or neurological insult. Participants then underwent neuropsychological assessment (Mini-International Neuropsychiatric Interview [MINI], inclusion cut-off =/< 2; the Folstein Mini Mental State Examination [MMSE], exclusion cut-off < 27; the Beck Depression Inventory [BDI-II], exclusion cut-off < 15; California Verbal Learning Task [CVLT-I English, CVLT-II French], exclusion cut-off based on recommendations by Norman, Evans, Miller, & Heaton, 2000). Only participants who met the above neuropsychological criteria and performed above chance on the practice context memory task presented in a Mock fMRI scanner were invited to return for a second visit and participate in the fMRI scanning portion of the study. All participants were paid for their participation, and the research ethics board of the Faculty of Medicine at McGill University approved the study protocol.

#### Visit 2: Task fMRI Session

##### Stimuli and Procedure

The task fMRI stimulus set has been used in previous studies and has been independently rated for pleasantness (Kwon et al., 2016; Rajah et al., 2010). Stimuli consisted of black-and-white photographs of faces that were varied in age and balanced for age and sex across experimental conditions. Each face presented during initial encoding was tested during subsequent retrieval, and participants were scanned during both encoding and retrieval memory phases (see Figure 1 for schematic representation of the task). A detailed description of the task paradigm used in the current study can be found in previous studies from our lab (Ankudowich et al., 2016, 2017).

**Figure 1:**
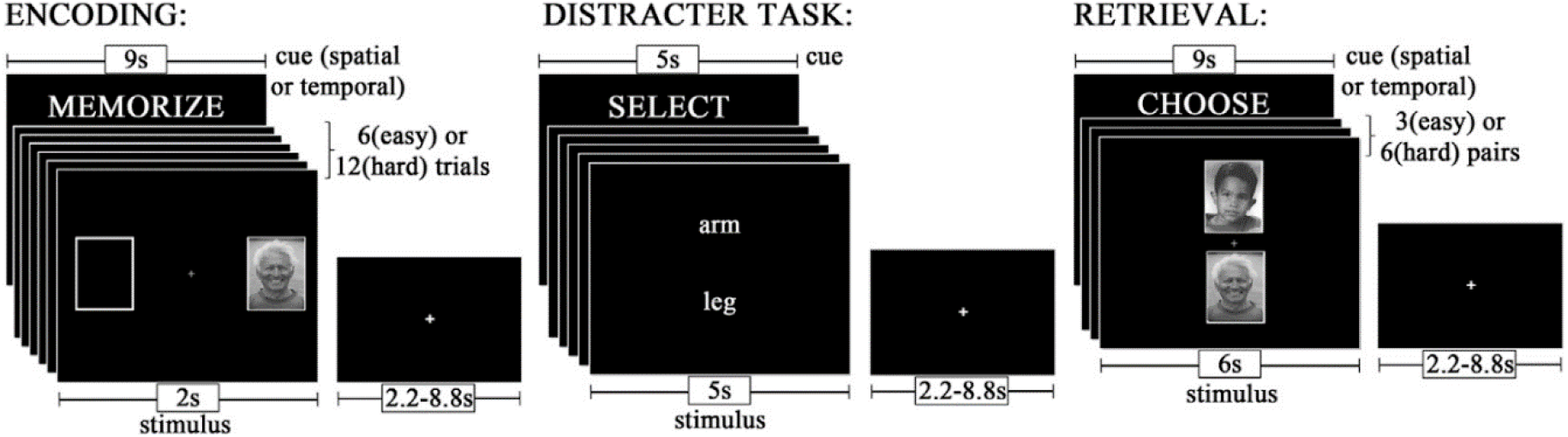
Task fMRI Procedure and event timeline.

Using a mixed rapid event-related design, participants were scanned across 12 experimental runs while they encoded and retrieved the spatial and temporal details of faces. Each run consisted of an ‘easy’ temporal context memory task (TE) and an ‘easy’ spatial context memory task (SE), and either a ‘hard’ temporal context memory task (TH) or a ‘hard’ spatial context memory task (SH). Easy and hard tasks differed in the number of stimuli to be encoded: 6 encoding stimuli for ‘easy’ tasks and 12 encoding stimuli for ‘hard’ tasks. In total, there were 72 trials presented for each encoding event type (i.e., 288 trials total) and 36 trials presented for each retrieval event type (i.e., 144 trials total). The current study focused on the behavioral and fMRI data collected during the spatial context memory tasks to compare our study findings with our previous activation analyses using the same paradigm (Subramaniapillai et al., 2019), and to further contextualize our work with the substantial psychological literature investigating sex differences in spatial episodic memory (Bender, Naveh-Benjamin, & Raz, 2010; De Frias et al., 2006; Gur & Gur, 2002; Herlitz et al., 1997; Sommer, Hildebrandt, Kunina-Habenicht, Schacht, & Wilhelm, 2013; Weiss et al., 2003; Yonker, Eriksson, Nilsson, & Herlitz, 2003; Young, Bellgowan, Bodurka, & Drevets, 2013). Our choice to only focus on the spatial context memory task further allows us to comprehensively address our aim of investigating sex differences in performance-related functional connectivity by comparing findings across several sex-aggregated and - disaggregated B-PLS analyses. Please refer to Ankudowich et al. 2016; 2017 for details regarding the temporal context memory tasks. Herein we present the details of the spatial context memory tasks.

Encoding was intentional, and at the start of each encoding phase, participants were cued (9 sec) to memorize the spatial location (whether a face appeared on the *LEFT* or the *RIGHT* during encoding) of the faces and to the level of task difficulty. At encoding, each face was presented (2 sec) on either the left or the right of a central fixation cross. There was a variable inter-trial interval (ITI) of 2.2 – 8.8 sec. During encoding, participants were instructed to rate the pleasantness of each face. Participants pressed a button with their right thumb to indicate a pleasant response and a button with their left thumb to indicate a neutral response using an MRI-compatible fiber optic response box. Between encoding and retrieval memory phases, participants performed a one-minute distractor task in which they were required to reverse alphabetize two words presented centrally on the computer screen. The distractor task was used to deter participants from actively rehearsing the encoding stimuli.

Following the distractor task, participants were presented with task instructions for retrieval (9 sec) to remind them of the spatial context task demands. During retrieval, participants were presented with pairs of previously encoded faces for 6 sec. One of the faces was presented above a central fixation cross, and the other was presented below. During the easy versions of the retrieval task, participants viewed 3 pairs of faces, and during the hard versions of the retrieval task, they viewed 6 pairs of faces. There was a variable ITI of 2.2 – 8.8 sec between retrieval events. For the spatial task, participants were asked to indicate which of the two faces was originally presented on the *LEFT/RIGHT*. Participants pressed a button under their right thumb to indicate a face at the top of the screen and they pressed a button under their left thumb to indicate a face at the bottom of the screen. Therefore, fMRI task-related activation for the spatial context memory paradigm was collected for four different event-types in this experiment: encoding spatial easy (eSE), encoding spatial hard (eSH), retrieval spatial easy (rSE), retrieval spatial hard (rSH).

##### Task fMRI Imaging Methods

Structural and functional magnetic resonance imaging data were collected at the Douglas Institute Brain Imaging Centre. Participants lied supine in a 3T Siemens Magnetom Trio scanner and wore a standard 12-channel head coil. T1-weighted anatomical images were first acquired for each participant at the start of the scanning session using a 3D gradient echo MPRAGE sequence (TR = 2300 msec, TE = 2.98 msec, flip angle = 9°, FOV = 256, 176 1 mm sagittal slices, 1 × 1 × 1 mm voxels). Blood-oxygen-level-dependent (BOLD) images were acquired with a single-shot T2*-weighted gradient echo-planar imaging (EPI) pulse sequence (TR = 2000 msec, TE = 30 msec, FOV = 256, matrix size = 64 × 64, in-plane resolution 4 × 4 mm, 32 oblique slices per whole-brain volume) while participants performed the context memory tasks. Visual task stimuli were back-projected onto a screen in the scanner bore using E-Prime software, and participants requiring correction for visual acuity wore plastic corrective lenses. A variable ITI (2.2 – 8.8 sec) was introduced to add jitter to event-related acquisitions.

#### fMRI Basic Preprocessing

Reconstructed images were preprocessed in SPM version 8 software. For each participant, the origin of functional images was reoriented to the anterior commissure of that individual’s acquired T1-weighted structural image. All functional images were then realigned to the first image, and motion artifacts were corrected using a 6-degree rigid-body transformation (three translation and three rotational parameters). Any experimental run in which within-run motion exceeded 1.5 mm was excluded from analysis. In total, 22 runs (1.2%) were excluded: 12 runs due to task noncompliance (e.g., failure to record participant responses, issues with the response box), 6 runs due to frontal/medial BOLD signal loss after fMRI preprocessing, 2 runs due to poor volumes, 2 runs due to scanner failure, and none due to excessive motion. Functional images were then normalized to an MNI EPI template and resliced at 4 × 4 × 4 mm voxel resolution and smoothed with an 8 mm full-width at half maximum (FWHM) isotropic Gaussian kernel. ArtRepair toolbox for SPM8 (http://cibsr.stanford.edu/tools/human-brain-project/artrepair-software.html) was used to correct slice artifacts prior to realignment and volume artifacts after normalization and smoothing (<5% interpolated data). Any run in which interpolated data exceeded 5% was excluded from analysis.

#### Analysis

##### Behavioral Data Analysis

###### Spatial Context Retrieval Accuracy and Reaction Time

Using R (R Core Team, 2013), we conducted robust linear mixed-effects regression (rlmer) models (using the robustlmm package; Koller, 2016) in the full cohort to test the three-way interaction between age, sex (2: men, women), and task difficulty (2: easy, hard) on retrieval accuracy (% correct) and reaction time (msec), respectively. The rlmer model is similar to the lmer model (Bates et al., 2015 for the lme4 package details) but additionally, it is robust to outliers by down-weighting the impact of extreme measures on the model performance (Koller, 2016). The models contained the random effect of participants to account for the variability of participants’ performance between the easy and hard versions of the spatial context task. The models used in terms of R syntax for spatial retrieval accuracy and reaction time, respectively, were:

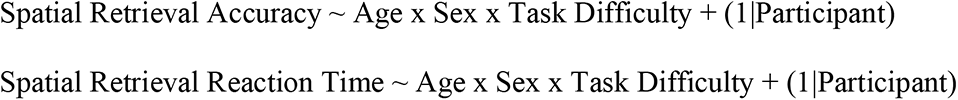

The continuous variable of age was standardized using a Z-score transformation, while the variables of sex and task difficulty were treated as categorical variables through deviation coding (−1, 1).

###### fMRI Preprocessing for PLS Connectivity Analysis Brain Parcellation

Figure 2 (below) illustrates the preprocessing steps used to generate the connectivity matrices for participants across the four task conditions, which were subsequently submitted to the PLS analysis. Using SPM’s MarsBaR toolbox, the average time series for 264 regions of interest (ROIs) defined by the Power et al. (2011) functional parcellation atlas were extracted for each subject for all task-related event-types across the full experiment. Each ROI was registered from the 2 x 2 x 2 mm^3^ Power et al. atlas to the 4 × 4 × 4 mm^3^ voxel resolution of our functional scans. To do this, we took each ROI’s central coordinates from the Power et al (2011) ROIs and identified a 7-voxel sphere surrounding the central coordinates. During this process of scaling down to the 4 x 4 x 4 mm^3^ voxel resolution, we eliminated ROIs with voxels that were not common to all participants and/or overlapped with other ROIs. We also excluded cerebellar ROIs because our fMRI acquisition did not completely acquire these regions, and the uncertain network ROIs because they did not belong to a major functional system in the brain. We additionally combined the memory retrieval network with the default mode network because the few nodes belonging to the memory retrieval network are activated in cognitive functions (e.g., memory, imagination) commonly attributed to the default mode network (Huo et al., 2018). Thus, we identified a total of 216 unique ROIs assigned to 9 brain networks: auditory, cingulo-opercular task control network (CON), default mode network (DMN), dorsal attention network (DAN), fronto-parietal task control network (FPN), salience, sensory/somatomotor network (SSM), visual attention network (VAN), visual (see Supplementary Table 1 for list of MNI coordinates and network affiliation).

**Figure 2.**
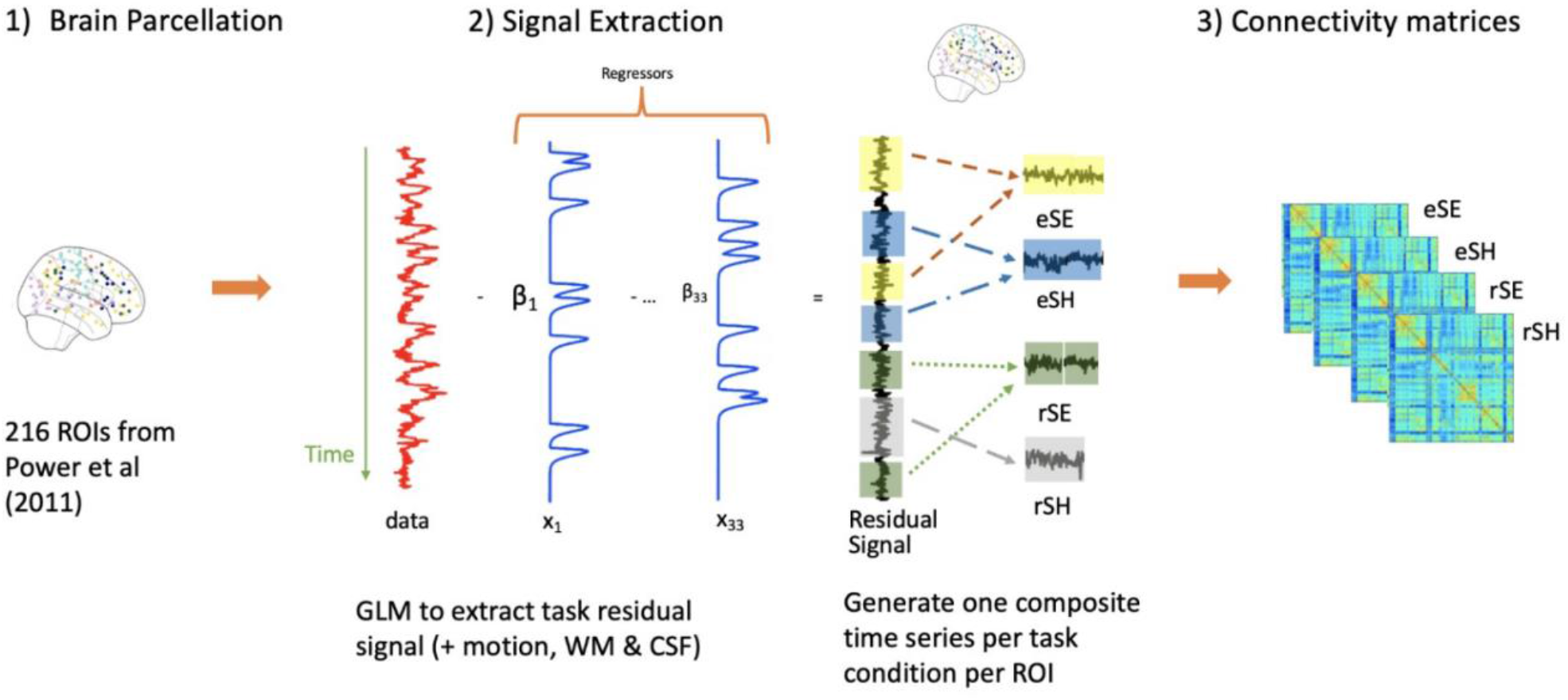
The fMRI preprocessing steps involved (1) functional parcellation of each subject across the 216 unique ROIs from the Power et al. atlas; (2) applying a GLM to extract the task residual signal after regressing 33 regressors to generate one composite time series per task condition for each ROI; (3) generating four connectivity matrices for each task condition for every participant. Note: ROI = region of interest, GLM = General Linear Model, WM = white matter, CSF = cerebrospinal fluid, eSE = encoding spatial easy, eSH = encoding spatial hard, rSE = retrieval spatial easy, rSH = retrieval spatial hard.

###### fMRI Signal Extraction

To examine task-related functional connectivity, it is recommended that first the mean task/event-related activity across the full experiment be regressed out of the fMRI signal. This accounts for the confound of task-timing-driven statistical associations (Cole et al., 2019). To this aim, event-related task activation for all 216 ROIs was estimated using SPM’s General Linear Model (GLM) with an ordinary least squares (OLS) approach (i.e., with AR(1) off), using a high pass filter set at 200 sec. This GLM consisted of 12 task-related regressors: correct subsequent memory events for all experimental tasks at encoding and retrieval, incorrect subsequent memory responses for all encoding tasks, incorrect context retrieval responses for all retrieval tasks, encoding and retrieval task instructions, and distraction task. In addition, the 6 movement regressors generated by SPM during motion correction, the mean white matter and the cerebrospinal fluid signals were also included as regressors in the GLM to correct for physiological noise (Birn et al., 2014). Finally, the temporal derivatives of the hemodynamic response function for each of the task-related regressors and the constant (i.e., intercept) resulted in a total of 33 regressors used in the GLM. Thus, this one GLM model was used to extract the mean residual time series for each ROI per event-type using the MarsBaR toolbox in SPM (http://marsbar.sourceforge.net/).

###### Generating Functional Connectivity Matrices

Since the focus of our current analysis is the spatial version of the task, we only generated functional connectivity matrices for each event-type of the spatial task. Each participant’s residual time series were concatenated across similar event-types to generate composite time series for each event-type. The minimum length of time for a concatenated event was 186 sec in the current study. Previous work has established that a minimum length of 30 sec is sufficient for reliable task-based connectivity analyses (e.g., Mohr et al., 2016). As a measure of functional connectivity, we computed Pearson correlations for each ROI with every other ROI across the time series. Connectivity matrices were created for each participant and event-type from the correlation coefficients, which then underwent Fisher z-transformation. Thus, in total, each subject had four connectivity matrices, one for each of the four event-types (i.e., eSE, eSH, rSE, rSH) of size 216 x 216. Since the matrix is symmetrical around the diagonal, there were a total of 23, 220 unique connections or edges.

###### PLS Functional Connectivity Analysis

Behavioral multivariate partial least squares (B-PLS) connectivity analysis was used to identify patterns of task-based functional connectivity (McIntosh & Misic, 2013), due to its ability to simultaneously detect distributed patterns of whole-brain connectivity that differ based on participants’ age, sex, and memory performance. We conducted two B-PLS connectivity analyses. The first was a **full group analysis (B-PLS1)**, in which we examined how age and memory performance in the full sample of adults (i.e., without disaggregating by sex) related to task-based connectivity during encoding and retrieval of SE and SH tasks. The second was a **between-sex (women, men) group B-PLS analysis (B-PLS2)**, in which we explored sex differences in age- and performance-related patterns of brain connectivity.

In the first analysis, connectivity matrices for each individual were organized by task event-type and then stored in a single group level fMRI connectivity matrix. In the second analysis, the between group factor of sex was included in the group level fMRI connectivity matrices. In both B-PLS analyses, normalized measures of participants’ age and retrieval accuracy were the behavioral measures of interest. We orthogonalized our behavioral vectors of age and accuracy to assess independent effects of age and performance (consistent with Subramaniapillai et al., 2019; see also Ankudowich et al., 2017). That is, prior to the PLS analyses, we conducted a regression analysis where task-specific retrieval accuracy was used to predict age to obtain an age-residual vector that would be uncorrelated with retrieval accuracy. These age-residual and retrieval accuracy vectors were then stacked in the same manner as the fMRI data matrix for each analysis, respectively (e.g., participant sex and by event-type for the between-sex group B-PLS). Given that the retrieval accuracy behavioral vector did not have age regressed from it, it allowed us to assess connectivity associated with age-related performance effects, whilst the age-residual allowed us to assess age effects orthogonal to performance effects. The following steps would be identical for both analyses, so they are presented once.

The stacked fMRI data matrix was then cross-correlated with the similarly stacked behavioral vectors. The resulting cross-correlation matrix was submitted to singular value decomposition (SVD). SVD re-expresses the matrix as a set of orthogonal singular vectors or latent variables (LV). Each LV consists of a singular value that reflects the proportion of matrix accounted for by that LV, and a pair of vectors (a left singular vector consisting of the behavioral weights and a right singular vector consisting of the connectivity weights) that reflect a symmetrical relationship between the pattern of whole-brain connectivity and the experimental design/behavior measures. The profile of behavioral weights shows how the behavioral vectors of age and retrieval accuracy are correlated to the pattern of whole-brain connectivity identified in the singular vector of connectivity weights. The connectivity weights identify the collection of edges that, as a group, are maximally related to the behavioral weights.

Significance testing for the LVs was done using 500 permutations (p < 0.05). The permutation test assesses whether the functional networks and behavioral profiles are more strongly associated with one another than expected by chance. Bootstrap resampling was performed to assess the reliability of each of the edges (500 bootstraps, bootstrap ratio [BSR] threshold was set at 95th percentile, p < 0.001). Connectivity edge contribution was estimated with edge loadings, which is calculated as the correlation of the participants’ PLS-derived brain score pattern with their stacked connectivity matrices. The pattern of edge loadings (i.e., correlations) is referred to as the loading matrix and reflects whether edges are more positively or negatively associated with the behavioral weights. A positive correlation coefficient in the loading matrix indicates a positive association with positive behavioral weights. Conversely, a negative correlation coefficient in the loading matrix is positively associated with the negative behavioral weights. Since the relationship between the behavioral weights and the loading matrix (i.e., connectivity weights) is symmetric, the inverse is also true. That is, a positive correlation coefficient indicates a negative association with negative behavioral weights and vice versa.

## Results

### Behavioral Results

Table 1 summarizes the participant demographic and neuropsychological information across the age groups for the full (n=141) and sex-disaggregated sample (n = 49 men, 92 women). Behaviorally, the rlmer model investigating the effects of age, sex, and task difficulty on memory accuracy showed a main effect of age (β = -0.03 [SE, 0.01]; t = -2.35, p < .05) and task difficulty (β = -0.04 [SE, 0.01]; t = -3.00, p < .05). Younger adults had greater accuracy than older adults on the tasks, and generally, participants performed worse on the SH task compared to the SE task. No other main effects or interactions were significant.

**Table 1.** Mean Demographic and Behavioral Measures (and Standard Errors)

|  | Total Behavioral<br>Sample | Total fMRI<br>Sample | Men | Women | *p* |
| --- | --- | --- | --- | --- | --- |
| Sample size (n) | 141 | 137 | 49 – Total<br>behavioral; 47 –<br>fMRI Sample | 92 – Total<br>behavioral; 90 fMRI<br>sample |  |
| Age (years) | 47.11 (1.41) | 47.26 (1.44) | 46.96 (2.44) | 47.20 (1.73) |  |
| Educations (years) | 15.73 (0.18) | 15.72 (0.18) | 16.06 (0.27) | 15.55 (0.23) | ns |
| Predicted full-scale IQ | 119.51 (0.44) | 119.60 (0.44) | 119.66 (0.73) | 119.43(0.56) | ns |
| BDI | 3.90 (0.32) * | 3.96 (0.32) * | 3.84 (0.53) | 3.93 (0.40) * | ns |
| CVLT-LFR | 13.17 (0.18) | 13.19 (0.19) | 12.35 (0.36) | 13.61 (0.19) | p<.05%# |
| CVLT-LCR | 13.43 (0.17) | 13.46 (0.17) | 12.76 (0.30) | 13.78 (0.20) | p<.05%# |
| CVLT-RG | 15.33 (0.69) | 15.36 (0.68) | 15.29(0.11) | 15.36 (0.09) | ns |
| BMI (kg/m<sup>2</sup>) | 24.26 (0.31) * | 24.25 (0.31) * | 24.49 (0.39) | 24.14 (0.43) * | p<.001^ |
| SE retrieval accuracy<br>(%correct) | 0.86 (0.01) | 0.86 (0.01) | 0.85 (0.01) | 0.86 (0.01) | p<.001# |
| SH retrieval accuracy<br>(%correct) | 0.83 (0.01) | 0.83 (0.01) | 0.80 (0.02) | 0.84 (0.01) | p<.001# |
| SE retrieval RT (msec) | 2474.95 (47.27) | 2488.80 (47.43) | 2417.32 (72.44) | 2505.31 (61.36) | p<.001# |
| SH retrieval RT (msec) | 2570.99<br>(43.85) | 2582.56<br>(43.94) | 2550.29<br>(72.25) | 2582.92<br>(55.35) | p<.001# |

\*One participant had missing information. Values in brackets represent the standard error. A linear regression of Age x Sex was performed on each of the measures (significance of $p < .05$ used) on the total sample (N=141). % The linear regression produced a significant effect of Sex, such that women outperformed men on this score. ^ Age x Sex interaction of BMI: age-related increase in BMI; younger and middle-aged adult men had higher BMI than their female counterparts; and older men had higher BMI than older women. # The linear regression produced a significant main effect of Age. The fMRI behavioral measures revealed that older adult participants performed significantly worse than younger and middle-aged participants and with significantly greater RT to complete the spatial tasks. BDI = Beck Depression Inventory; LFR = Long-form Free Recall; LCR = Long-form Cued Recall; RG = Recognition; BMI = body mass index.

There were also significant main effects of age (β = 145.60 [SE = 68.71]; t = 2.12, p < .05) and task difficulty (β = 130.23, [SE = 36.71]; t = 3.55, p < .05) on reaction time. Young adults were faster than older adults across SE and SH tasks, and participants took longer to respond to the SH task than the SE task. No other main effects or interactions were significant. Therefore, there were no sex differences, nor sex-by-age interactions in task performance.

### Functional connectivity results

Four participants’ fMRI images failed preprocessing and were excluded from the PLS analyses (2 women and 2 men). Therefore, the sample size for the PLS analyses was 137 (47 men and 90 women). Figures 2 through 5 depict the relevant information for the significant LVs in both the full group B-PLS1 and the between-sex group B-PLS2 analyses, respectively. The subplots include the 1) thresholded loading matrix, 2) behavioral correlation weights, 3) network density matrix, and 4) brain figure representing the highly involved nodes. The thresholded connectivity matrix (1) represents the 95th percentile of the z-score values of correlation coefficients. The behavioral weights (2) indicate how the loading matrix relates to the behavioral vectors of age and accuracy in women and men. The network density matrix (3) represents the sum of the unthresholded significant edges divided by the total number of possible edges between any two networks (or within a network). Each LV generated two density plots because calculations were done separately on the positive and negative correlation coefficients. Density matrices that produced sparse significant edges (<5%) were not included. Finally, the brain figures (4) identify the most highly contributing nodes from the thresholded loading matrix, as determined by the ranked sum of the correlation values from most to least involved. Below we report the detailed findings of each B-PLS analysis.

#### Full Group B-PLS1 Results

The full group B-PLS1 analysis examining age and performance effects in connectivity identified two significant LVs at p < 0.05. The first LV (LV1, accounting for 70.15% cross-block covariance) identified significant positive connectivity weights (in red) between several networks (Figure 3A).

**Figure 3.**
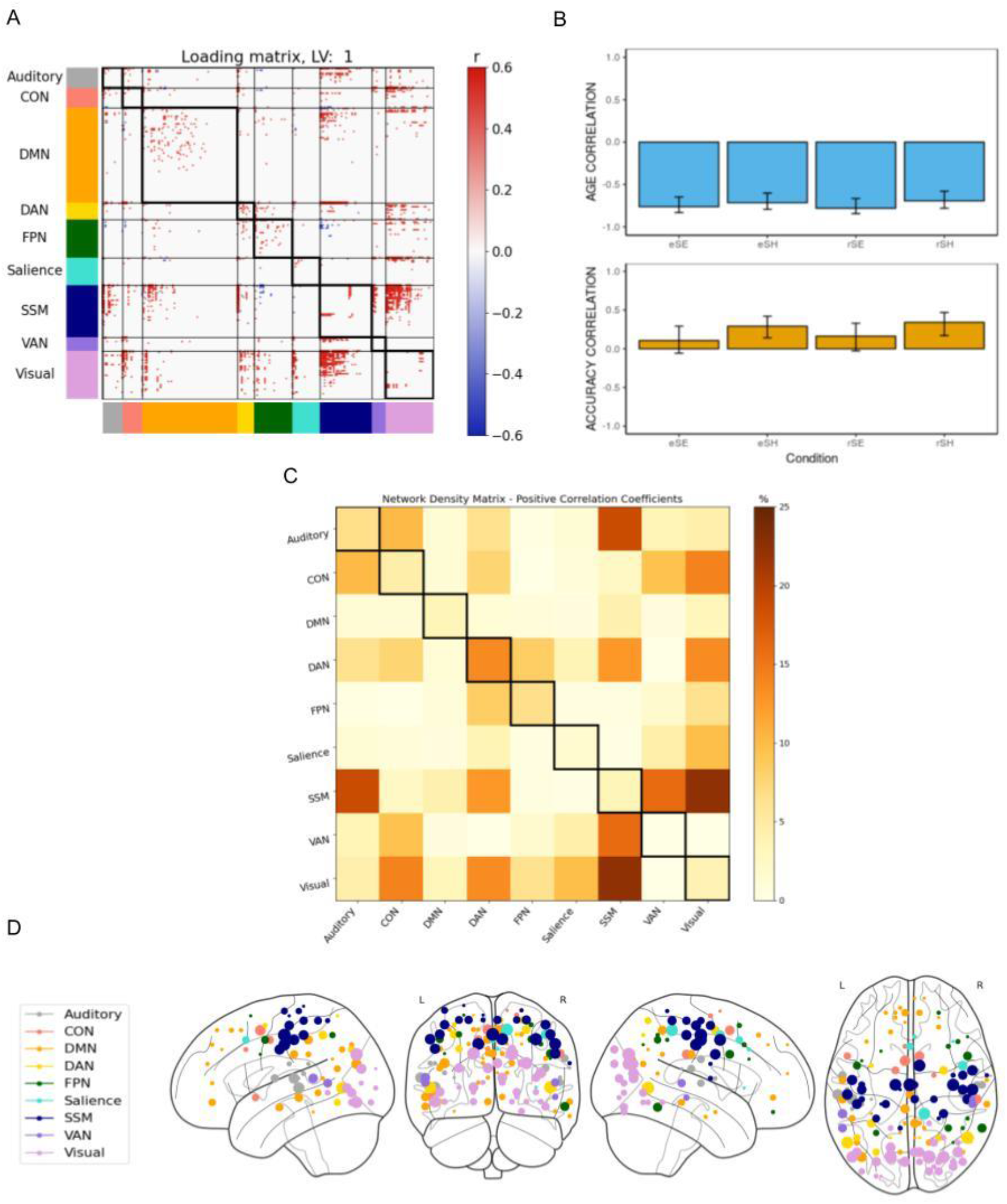
B-PLS1, LV1: Differential effects of age & accuracy on task-related brain connectivity. B-PLS1, LV1 reflects differences in how age and accuracy on the task influence task-related brain connectivity. **(A)** Thresholded 95th percentile of correlations between participants’ task fMRI data and behavioral profile indicated in B. **(B)** Correlation between the behavioral vectors of age and accuracy with the task fMRI connectivity of participants (behavior correlation weights). Error bars represent bootstrapped standard deviations. **(C)** The density plot for the positive correlation coefficients (i.e., sum of the significant correlation coefficients after thresholding, divided by the total number of edges between any two networks). The density matrix for the negative correlation coefficients is not presented because there were no significant edges. **(D)** Most densely connected nodes from the positive salience loading matrix as represented by the rank sum of the correlation coefficients of the thresholded matrix. Greater node size represents greater node involvement. eSE = encoding spatial easy; eSH = encoding spatial hard; rSE = retrieval spatial easy; rSH = retrieval spatial hard; CON = cingulo-opercular network; DMN = default mode network; DAN = dorsal attention network; FPN = frontoparietal network; SSM = somatomotor network; VAN = ventral attention network.

The loading matrix and density matrix for LV1 (Figures 3A and 3C) indicates that there were three dominant patterns of positive connectivity involving the DAN, visual network, and SSM network. First, LV1 identified positive within-network connectivity weights in the DAN and FPN, and between the DAN and FPN, SSM, and visual network. Second, there was positive network connectivity between the (i) visual network and CON, and (ii) SSM and the auditory network and VAN. The matrices and behavioral correlation weights (Figure 3B) together indicates that this pattern of positive brain connectivity was negatively correlated with age across all encoding and retrieval conditions and was positively correlated with memory performance during the hard spatial context memory task. Specifically, greater positive functional connectivity among these networks during the encoding and retrieval phases of the hard, but not easy, spatial context memory task was positively correlated with memory accuracy but negatively correlated with age. Therefore, LV1 identified patterns of task-related functional connectivity that differentiated age and memory performance effects for the hard spatial context memory tasks.

The second LV accounted for 17.47% cross-block covariance and identified only significant negative connectivity weights (in blue) as seen in the loading matrix (Figure 4A). The density matrix (Figure 4C) identified dense patterns of connectivity between DAN and auditory, CON, DMN and VAN. Taken together with the behavior correlation weights (Figure 4B), these networks showed a negative correlation with retrieval accuracy. That is, greater connectivity between these networks during encoding and retrieval was related to poorer performance for all memory tasks.

**Figure 4.**
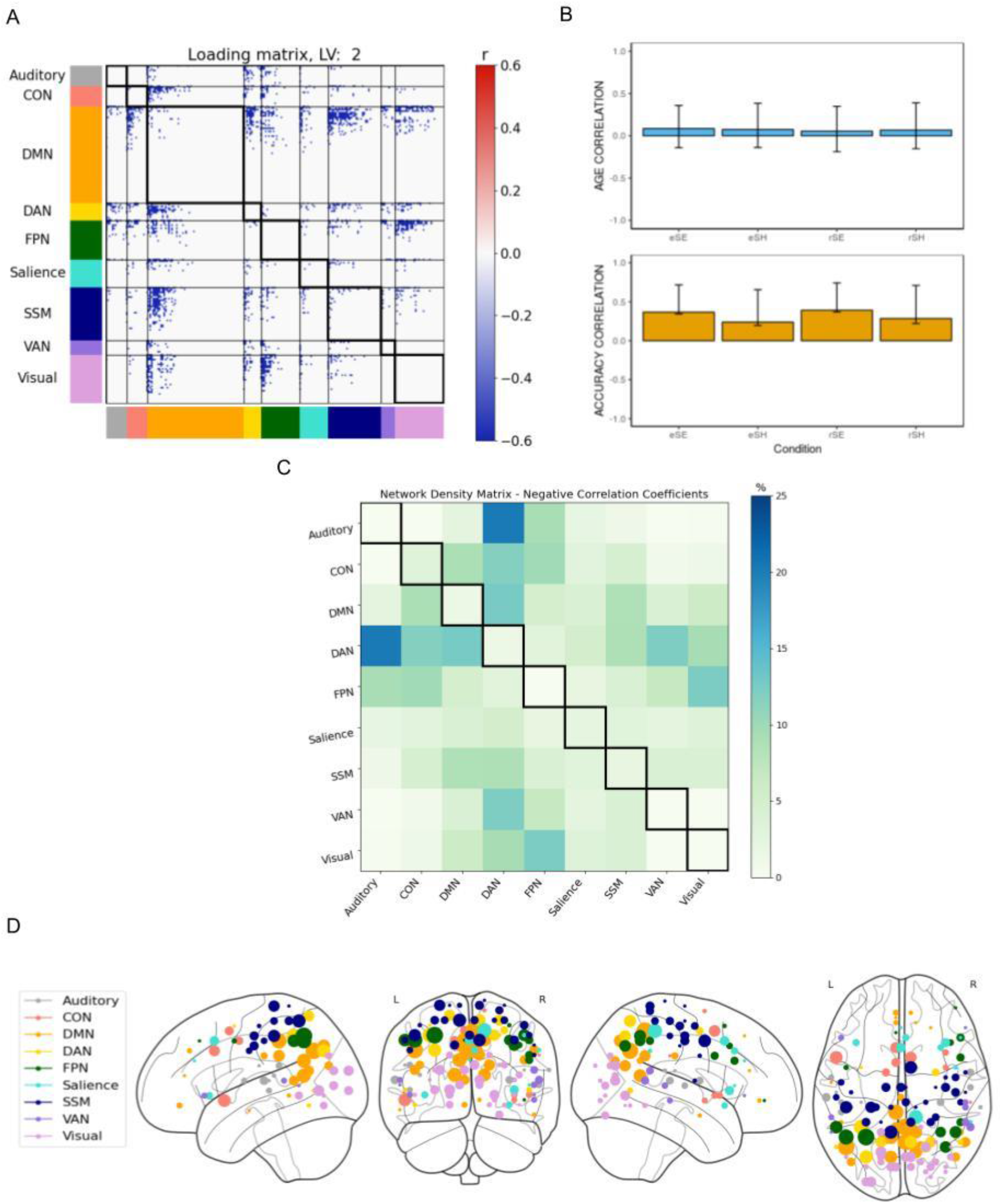
B-PLS1, LV2: Accuracy- but not age-related effects on task-related brain connectivity. B-PLS1, LV2 reflects how accuracy was related to task-related brain connectivity but not age. **(A)** Thresholded 95th percentile of correlations between participants’ task fMRI data and behavioral profile indicated in B. **(B)** Correlation between the behavioral vectors of age and accuracy with the task fMRI connectivity of participants (behavioral correlation weights). Error bars represent bootstrapped standard deviations. **(C)** The density plot for the negative correlation coefficients (i.e., sum of the significant correlation coefficients after thresholding, divided by the total number of edges between any two networks). The density matrix for the positive correlation coefficients is not presented because there were no significant edges. **(D)** Most densely connected nodes from the negative salience loading matrix as represented by the rank sum of the correlation coefficients of the thresholded matrix. Greater node size represents greater node involvement. eSE = encoding spatial easy; eSH = encoding spatial hard; rSE = retrieval spatial easy; rSH = retrieval spatial hard; CON = cingulo-opercular network; DMN = default mode network; DAN = dorsal attention network; FPN = frontoparietal network; SSM = somatomotor network; VAN = ventral attention network.

#### Between-Sex Group B-PLS2 Results

The between-sex group B-PLS2 analysis examining age and performance effects separately in women and men identified four significant LVs at p < 0.05. Since LV1 and LV2 accounted for most of the original variance in data (87.62%), we present and discuss the findings for LV1 and LV2 as they would represent the most valuable information with regards to sex differences in age and memory accuracy on task-related functional connectivity (Zeng & Wang, 2010). The results and figures for LV3 and LV4 are reported in the Supplementary Figures 1 and 2. The results and figures for LV3 and LV4 are reported in the Supplementary Figures 1 and 2.

LV1 accounted for 44.58% of cross-block covariance and showed both significant positive and negative connectivity weights. The behavior correlation plot indicates that the patterns of connectivity identified by LV1 was differentially correlated with age and memory performance during hard spatial context memory tasks in men and women, recapitulating the LV1 effect of the full group B-PLS1. The loading and density matrices (Figure 5A, C, D) showed dense positive connections involving DAN, SSM, and visual networks, consistent with LV1 from the B-PLS1. However, by disaggregating our connectivity analysis by sex we observed that the positive functional connectivity patterns also support retrieval performance during easy spatial context memory tasks in women only (i.e., the confidence interval does not contain zero). Furthermore, a unique pattern of negative weighted connectivity involving CON, DAN, FPN, and SSM was also identified. In both sexes, age was positively correlated with increased connectivity between SSM and DAN, FPN and between CON and FPN, while memory performance during hard spatial context memory tasks was negatively correlated with this pattern of connectivity in both sexes, and during easy spatial context retrieval in women only.

**Figure 5.**
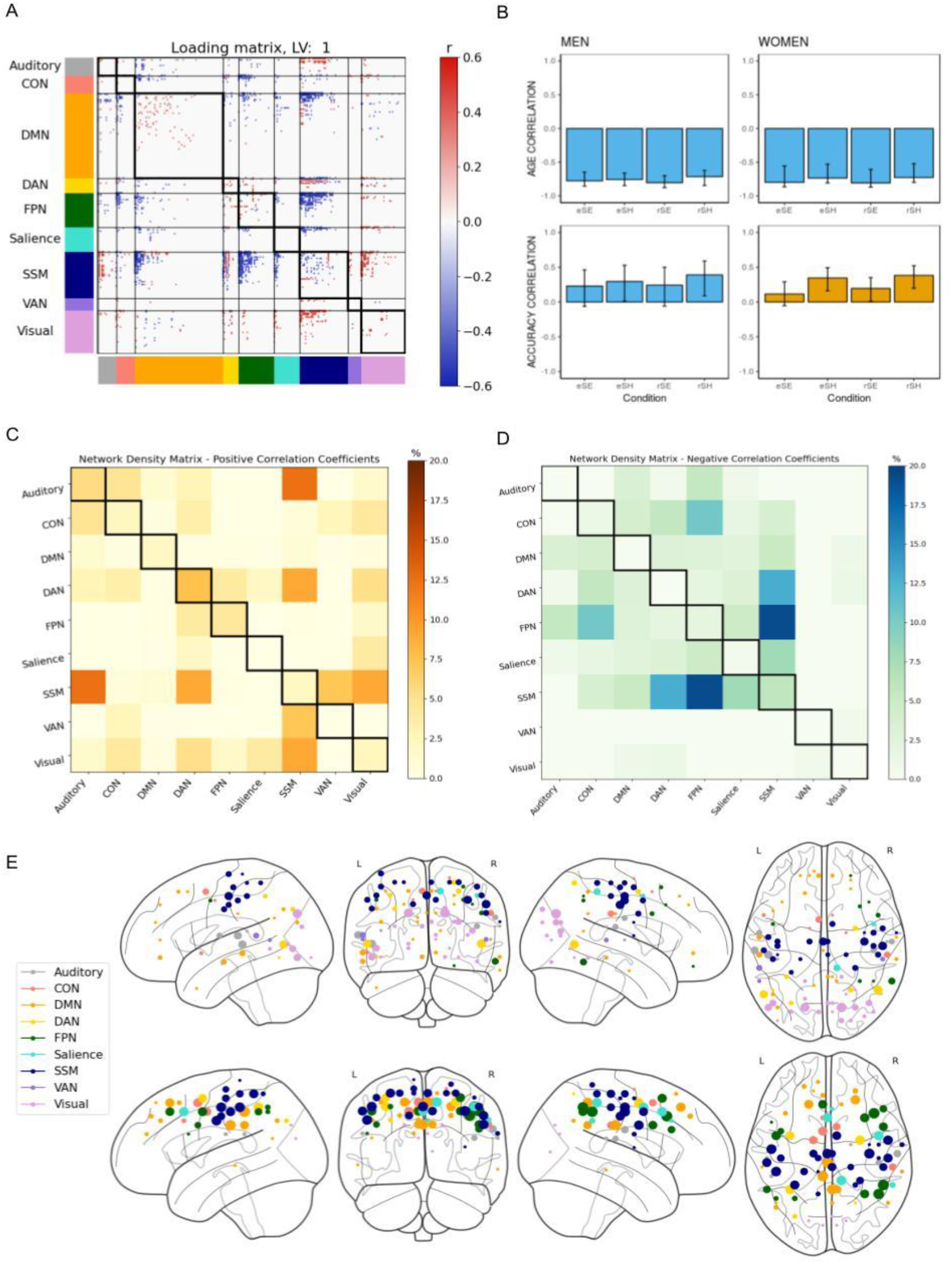
B-PLS2, LV1: Sex similarities in age and accuracy effects on task-related brain connectivity. B-PLS2, LV1 sex similarities in age and performance on task-related brain connectivity. **(A)** Thresholded 95th percentile of correlations between participants’ task fMRI data and behavioral profile indicated in B. **(B)** Correlation between the behavioral vectors of age and accuracy with the task fMRI connectivity of participants (behavioral correlation weights). Error bars represent bootstrapped standard deviations. **(C)** The density plot for the positive correlation coefficients (i.e., sum of the significant correlation coefficients after thresholding, divided by the total number of edges between any two networks). **(D)** The density plot for the negative correlation coefficients. **(E)** Most densely connected nodes from the positive (top) and the negative (bottom) salience loading matrix as represented by the rank sum of the correlation coefficients of the thresholded matrix. Greater node size represents greater node involvement. eSE = encoding spatial easy; eSH = encoding spatial hard; rSE = retrieval spatial easy; rSH = retrieval spatial hard; CON = cingulo-opercular network; DMN = default mode network; DAN = dorsal attention network; FPN = frontoparietal network; SSM = somatomotor network; VAN = ventral attention network.

LV2 accounted for 21.66% of the cross-block covariance and identified significant positive between-network connections involving DAN, SSM and the visual network (Figure 6A and 6C). The behavior correlation weights (Figure 6B) indicates there were sex differences in how age and memory performance correlated with this pattern of task-related brain connectivity. In men, positive connectivity among these networks was negatively correlated with memory performance across all tasks; and age was related to increased connectivity among these networks only during easy spatial context memory tasks. In contrast, in women, memory performance was not related to connectivity among these networks, but age was negatively correlated with connectivity in these networks across all tasks. Therefore, LV2 identified sex differences in how both age and memory performance correlated with task-based brain connectivity.

**Figure 6.**
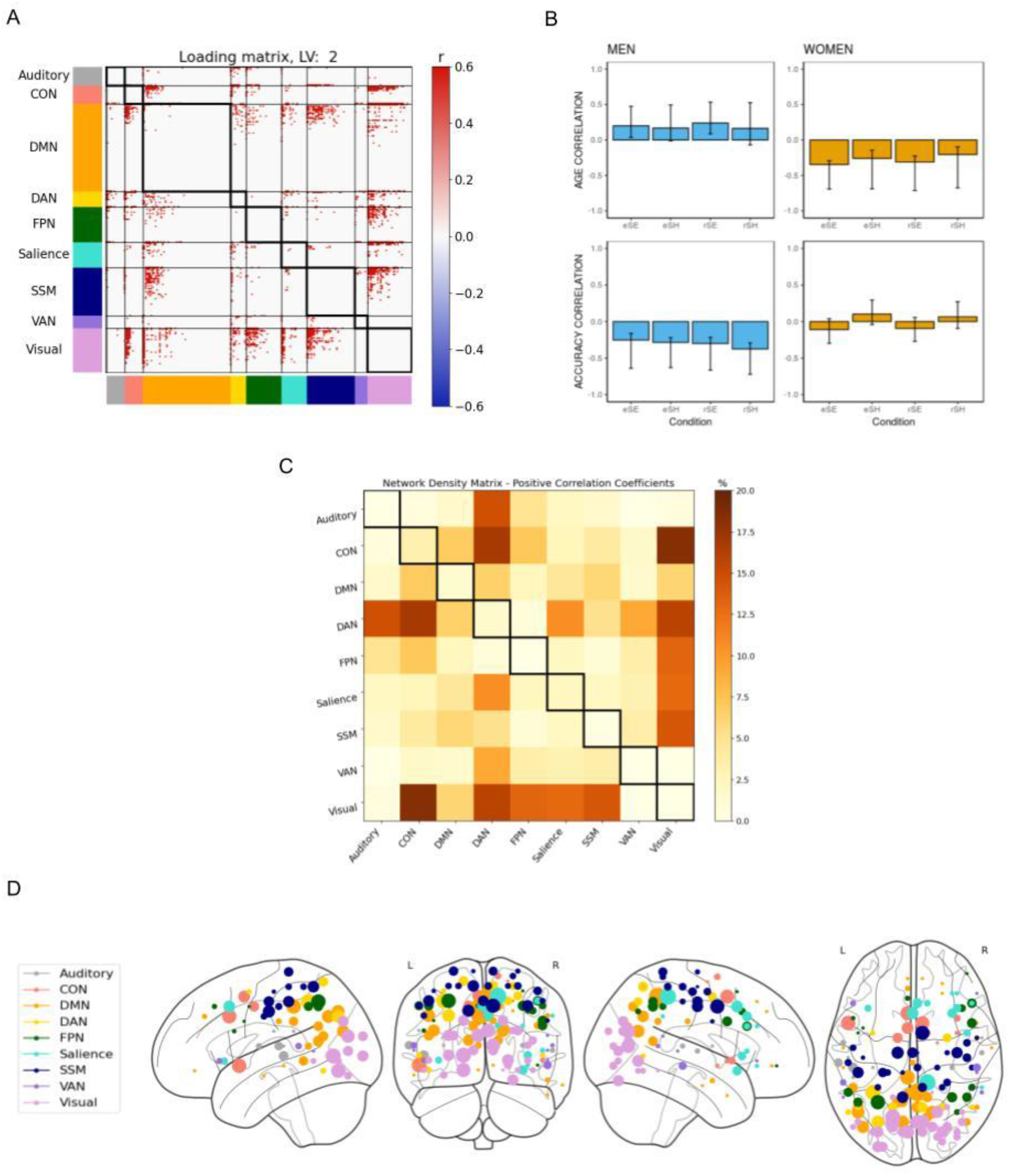
B-PLS2, LV2: Sex differences in age and accuracy effects on task-related brain connectivity. B-PLS2, LV2 sex differences in age and performance on task-related brain connectivity. **(A)** Thresholded 95th percentile of correlations between participants’ task fMRI data and behavioral profile indicated in B. **(B)** Correlation between the behavioral vectors of age and accuracy with the task fMRI connectivity of participants (behavioral correlation weights). Error bars represent bootstrapped standard deviations. **(C)** The density plot for the positive correlation coefficients (i.e., sum of the significant correlation coefficients after thresholding, divided by the total number of edges between any two networks). **(D)** Most densely connected nodes from the positive salience loading matrix as represented by the rank sum of the correlation coefficients of the thresholded matrix. Greater node size represents greater node involvement. eSE = encoding spatial easy; eSH = encoding spatial hard; rSE = retrieval spatial easy; rSH = retrieval spatial hard; CON = cingulo-opercular network; DMN = default mode network; DAN = dorsal attention network; FPN = frontoparietal network; SSM = somatomotor network; VAN = ventral attention network.

### Supplementary Analyses

We performed several post-hoc analyses to account for confounding factors that may have influenced the findings and subsequent interpretation of our primary analyses. First, sex differences in education and intracranial volume (ICV) may have impacted our study findings. Men typically have larger ICV than women (Ruigrok et al., 2014) and education level may have a strong involvement as a gendered reserve contributor (Subramaniapillai et al., 2021). Thus, we ran a supplementary analysis using a sub-cohort (n = 48) of women and men selected from our full sample matched according to age, education, and ICV to determine whether the LV patterns identified in our primary analyses were similar after controlling for these factors. This supplementary analysis revealed similar findings as those presented in our primary analyses (results presented in Supplementary Figures 3 and 4).

Second, while the choice to regress mean task-related activity is grounded in previous literature (Cole et al., 2019), we conducted supplementary B-PLS analyses without regressing mean task-related activity to enable readers to compare findings across differences in this preprocessing methodology. The LV effects from this supplementary analysis were consistent with our primary analysis (Supplementary Figures 5 and 6).

## Discussion

The goals of the current study were two-fold. First, we used behavior partial least squares (PLS) connectivity analysis to test the hypothesis that age and memory performance (retrieval accuracy) would be inversely associated with task-based connectivity between the DAN, DMN and FPN during successful encoding and retrieval of face-location associations (spatial context memory). We then disaggregated our analyses by self-reported sex and tested the hypothesis that there would largely be similarities in performance-related connectivity in both sexes and sex differences in the effect of age on memory performance-related brain connectivity, consistent with our prior task-based activation analyses of sex differences during spatial context memory (Subramaniapillai et al., 2019). The behavioral data from the current study replicated our prior work based on smaller sample sizes: there was no significant effect of sex on accuracy and reaction time, nor any significant interactions of age and sex. There were significant main effects of age and task difficulty on spatial context memory accuracy and reaction time, as reported previously (Ankudowich et al., 2017; Subramaniapillai et al., 2019).

The multivariate behavior PLS results from the full group B-PLS1 and between-sex group (women, men) B-PLS2 results generally corroborated our age-related hypotheses. Age and memory performance were inversely correlated to connectivity between DAN, FPN and visual networks in both sexes. Aging was also related to greater between-network integration among non-sensory networks, which was related to lower performance on hard spatial context memory tasks in both sexes, and lower performance during easy spatial context retrieval in women only. However, our sex-related hypotheses were not supported. We observed both similarities and differences in age-related and performance-related patterns of task-based functional connectivity, which did not differ by memory phase (encoding and retrieval). We discuss the details of our connectivity results below and highlight the importance of disaggregating task-based connectivity results by sex and gender in computational and clinical neuroscience studies of normative aging and episodic memory function.

### Sex similarities in age- vs. performance-related patterns of task-based connectivity during spatial context memory encoding and retrieval

In both B-PLS analyses, LV1 indicated that in both women and men, better memory performance during hard spatial context memory tasks was related to increased positive connectivity: (i) between DAN and the FPN, SSM, and visual networks, (ii) between SSM and the VAN, auditory, and visual networks, and (iii) within the DAN and FPN during encoding and retrieval phases of the hard spatial context memory tasks. In contrast, age was associated with decreased connectivity among these networks across all task conditions in both sexes (B-PLS1, LV1 and B-PLS2, LV1). This pattern of connectivity was correlated only with memory performance during hard but not easy tasks, which suggests increasing encoding load and retrieval demands during the spatial context hard > easy tasks, resulting in the engagement of several domain-general cognitive control and attention-related brain networks (i.e., DAN, FPN) to support memory performance. This observation is consistent with prior brain activation studies that have highlighted the importance of attention and cognitive control processes for successful episodic encoding and retrieval (Ciaramelli & Moscovitch, 2020; Smallwood et al., 2021), particularly for the memory of source and/or contextual details (Dulas & Duarte, 2014; Rajah, Ames, & D’Esposito, 2008; Rajah et al., 2010; Thakral, Wang, & Rugg, 2015). Also, we observed that across encoding and retrieval, men and women exhibited similarities in performance-related functional connectivity. This indicates that successful memory performance during the hard spatial context tasks relied on the reinstatement of functional connections present at encoding, during the later retrieval phase. This finding is consistent with current theories emphasizing the importance of recapitulation of cognitive/brain states and episodic replay to support retrieval success (Hill, King, & Rugg, 2021; Morcom, 2014; Stawarczyk, Wahlheim, Etzel, Snyder, & Zacks, 2020; Wimmer, Liu, Vehar, Behrens, & Dolan, 2020). Moreover, our current findings indicate this reinstatement occurs at a broad network level and is associated with individual differences in retrieval success. The finding that greater DAN-FPN connectivity during encoding and retrieval was correlated with better performance during harder spatial context memory tasks and younger age is consistent with prior studies that reported that FPN connectivity with DAN supports episodic memory, and with our hypothesis that age-related declines in episodic memory are related to reduced DAN-FPN connectivity (Avelar-Pereira et al., 2017; Benoit & Schacter, 2015; Cabeza & St Jacques, 2007; Habeck et al., 2012; Kim, 2012; Spreng et al., 2016). Beyond these predicted results, our task fMRI connectivity results highlight that the distinct pattern of connectivity among the visual network, SSM, and higher-order CON and DAN networks supported successful encoding and retrieval during hard spatial context memory in both women and men, and easy spatial context retrieval in women. Greater sensory and SSM connectivity in both sexes likely reflected the complex sensory-motor remapping demands of the task. At encoding, stimuli were presented left/right; at retrieval, two old faces were oriented top/bottom, but retrieval was based on a left/right decision and response data were collected from a horizontally oriented response box. The vertical presentation at retrieval was done to avoid stimulus masking effects, however, likely increased the stimulus-response mapping demands of the spatial context memory task (Power et al., 2011). Thus, age-related decreases in these connectivity patterns may reflect reductions in the ability to attend and integrate visual and sensorimotor information with goal-directed cognitive control processes. This may in turn have contributed to poorer memory function in both women and men. The observation that this pattern of connectivity was only correlated with better performance on hard tasks in both sexes is consistent with prior studies showing modulation of frontoparietal cognitive control processes as a function of task difficulty across cognitive tasks, including episodic memory tasks (Cole & Schneider, 2007; Dobbins & Han, 2006; Kim, 2010; Rajah et al., 2008; Rajah et al., 2011; Vincent et al., 2008). Interestingly, in women the correlation between connectivity and memory performance was also observed for easy spatial context retrieval and points to a sex difference in task-related functional connectivity that is discussed in greater detail below.

### Sex differences in the performance-related task-based connectivity during easy spatial context retrieval

The full group and between-sex group PLS LV1 results supported the hypothesis that aging in women and men was related to declines in within-network segregation in DAN and FPN. However, only after disaggregating our analysis by sex did we observe the predicted age-related increase in between-network connectivity (integration) among non-sensory networks, *i.e.,* CON, DMN, DAN, FPN, salience and SSM, across all task conditions in both women and men (B-PLS2, LV1, negative connectivity matrix).This pattern of connectivity was negatively correlated with memory performance during hard spatial context memory tasks in both sexes, and with memory performance during easy spatial context memory tasks in women only. Therefore, by disaggregating our analyses by sex, we were able to identify sex differences in performance effects related to easy spatial context retrieval.

This result indicates that the age effects identified in LV1 had a more general effect on memory performance in older, compared to younger women; but only affected memory performance on hard spatial context memory tasks in older, compared to younger men. Moreover, it is possible that the between-network integration observed in the sex disaggregated, but not the full group, analyses, may have been driven by performance effects in older women during the easy spatial context retrieval conditions. We have previously observed greater generalization in activation patterns across women, compared to men, in the activation analysis of a smaller sample of adults who participated in the current study (Subramaniapillai et al., 2019) and in a sample of older adults with a family history of late-onset AD (Rabipour et al, 2021). The current results shows that greater between-network integration was apparent at both levels of task difficulty in women only and may reflect increased generalization (or dedifferentiation) of function as women age (Chan et al., 2014).

### Sex differences in age- and performance-related patterns of task connectivity

Based on prior resting state fMRI connectivity studies (Avelar-Pereira et al., 2017; Ferreira et al., 2016; Jockwitz et al., 2017; Klaassens et al., 2017; Spreng & Schacter, 2012; Zonneveld et al., 2019), we hypothesized that there would be age-related increases in DAN-DMN task-based connectivity during encoding and retrieval, which would be inversely correlated with memory performance. Both our full group B-PLS1, LV2 and between-sex group B-PLS2, LV2 indicated that increased connectivity between DAN-DMN during spatial context encoding and retrieval was related to poorer memory performance. However, it was only after we disaggregated our analysis by sex, we observed the predicted age effect – and only in men. Specifically, men showed age-related increases in DAN-DMN connectivity during easy spatial context memory encoding and retrieval tasks, which was negatively correlated to their memory performance. Men also exhibited weak connectivity between DAN-FPN and an increased connectivity pattern between DMN and the auditory, CON, and visual networks. This suggests that decoupling of DAN-FPN, greater DAN-DMN connectivity, and greater connectivity between DAN and FPN with sensory networks was correlated with men’s poorer episodic encoding and retrieval. This result is consistent with the hypothesis that suppression of DAN-DMN connectivity and increased DAN-FPN connectivity during externally oriented tasks, i.e., episodic memory tasks, supports successful task performance (Smallwood et al., 2021; Spreng & Turner, 2019), but highlights that this age-related deficit in the suppression of DAN-DMN connectivity was specific to men in the current study. Furthermore, these age- and performance-related differences in connectivity in men, suggests they may exhibit decreases in top-down attentional control of visual processing with age that was detrimental to performance (Esposito et al., 2018; Grady et al., 2016; Vogel et al., 2012). This is also consistent with our prior activation analysis demonstrating that with advanced age, men engaged visual sensory processing areas for successful memory performance, possibly relying on task strategies related to semantic processing (Subramaniapillai et al., 2019).

Women, in contrast, exhibited an age-related decrease in DAN-DMN connectivity and in DAN connectivity with other networks. Moreover, this age-related difference in connectivity was not related to memory performance in women. Thus, age-related memory decline in women in the current study was not associated with altered DAN-DMN connectivity. This was contrary to our hypothesis that similar age effects would be observed in women and men, and indicates that in women, age-related spatial context memory decline was primarily represented by the effects observed in B-PLS2 LV1 (discussed above). More broadly, our findings indicate there were sex differences in DMN and DAN connectivity with age. This may be indicative of different task orientations in older women, compared to men (Ankudowich et al., 2017); or reflect sex differences in the rate at which age effects functional connectivity (Scheinost et al., 2015). Indeed, using resting state functional connectivity, Scheinost et al (2015) reported that between the ages of 18 and 65 yrs of age, men exhibited steeper differences in DMN connectivity by decade, compared to women. Given the fact that age-related cognitive decline and neurodegenerative diseases, i.e., AD has been linked to altered connectivity involving the DMN (Hafkemeijer, van der Grond, & Rombouts, 2012), future work should further explore if there are sex differences in task-based DMN connectivity in other memory paradigms, and at rest.

#### Caveats

The present study examined sex similarities and differences in spatial context memory across the lifespan using a novel functional connectivity methodological approach. However, our study has several limitations that future work should address. First, our findings are specific to the tasks analyzed and future studies aimed at replicating results in different episodic memory paradigms is essential to validating the generalizability of our current finding. Second, a comprehensive data collection approach was not used when collecting participants’ biological sex or menopause status. Our current study acquired participants’ biological sex through self-report, although it could also be ascertained through other means, including participants’ sex hormone measurements. Hormone collection is especially relevant when investigating major life transitions, such as menopause, which is associated with age-related differences in women’s hormonal profiles. As a consequence of women’s greater menopause-related hormonal changes and the established literature of memory effects during this transition (Henderson, 2010; Li, Cui, & Shen, 2014; Rentz et al., 2017; J. Yonker et al., 2006), we decided to omit our cohort of women transitioning through menopause and those who underwent HRT. Although our small cohort size of women in the menopause transition prevented us from including them in our primary analysis, it is essential that future research integrate important life transitions to better inform our understanding of healthy aging models in women and men. Lastly, given that we did not collect information about participants’ sociocultural gender, it is further challenging to disentangle the effects of biological sex and sociocultural gender on age- and performance-related connectivity differences.

Also, our relatively small cohort size constitutes another limitation of the current study. Despite the small cohort, our findings complement our previous activation studies, both at the behavioral and functional level, using the same lifespan cohort (Subramaniapillai et al., 2019; Ankudowich et al., 2016; 2017). Moreover, we found that our PLS connectivity findings were robust to several methodological confounds. First, one challenge that we foresaw was that sex differences in intracranial volume (ICV), with men typically having greater ICV than women, may be driving our functional connectivity results. However, when we ran our analysis on a smaller cohort of participants matched on ICV (and age and education), our findings corroborate our primary analysis (Supplementary Figures 3 and 4).

Finally, although we have theoretical justification for regressing task mean activity from the fMRI signal, one might rightfully ask what the error term actually means, in terms of functional relevance. When we ran the PLS connectivity analysis without regressing mean task-related activity, the analysis generated the same exact LV results and functional network connectivity with minimal differences observed in connectivity at the nodal rather than network level. This enabled us to conclude that the level of interpretation we used for the current study (i.e., at the network level) would have resulted in the same interpretations of findings, whether or not we chose to regress mean task-related activity. Future work should endeavour to understand what these minute differences mean at the node level, both theoretically and conceptually. Thus, although there was the possibility of several confounds, our supplementary analyses findings demonstrate our primary analysis was robust to different preprocessing strategies and methodological confounds.

## Conclusions

The current study is the first to examine age- and performance-related differences in task-based connectivity during episodic encoding and retrieval in a normative adult lifespan sample, and to explore how self-reported sex effects these patterns of connectivity. In both sexes, age- and memory performance were inversely correlated with DAN-FPN connectivity. In addition, we observed the predicted age-related increase in DAN-DMN connectivity but only in men, while women showed more between-network integration and generalization of function with advanced age. Thus, different neurocognitive mechanisms contribute to normative age-related differences in episodic memory in women and men. These sex and gender differences should be considered when interpreting task-related and resting-state fMRI studies of AD, and other age-related neurological and psychiatric diseases that have sex differences in prevalence rates and are known to affect individuals’ episodic memory function (i.e., Parkinson’s disease). Overall, our results highlight the importance of considering sex and gender in study design, analysis, and interpretation in cognitive neuroscience studies of aging and memory.

## Acknowledgements

We thank all the research participants who made this work possible. This work was supported by CIHR Operating Grants (GS9-171369 and 201610PJT-374992) and NSERC Discovery Grant (RGPIN-2018-05761) awarded to M.N. Rajah; Canada Research Chair II to B. Misic; the Natural Science and Engineering Research Council Graham Bell Canada Graduate Scholarship-Doctoral and the Healthy Brains Healthy Lives Doctoral Fellowship awarded to S. Subramaniapillai.

## Contributions

M. N. Rajah (MNR) designed the study. S. Subramaniapillai (SS), S. Rajagopal (SR) and E. Ankudowich (EA) contributed to data processing and analysis. S. Pasvanis (SP) and EA led data collection and quality control. SS and SR created figures and tables. Bratislav Misic (BM) provided the PLS connectivity code, SR edited and created the GitHub code used in the current publication. SS, SR, EA, BM, and MNR provided analytic, theoretical input and editorial feedback on drafts of this paper. EA wrote and earlier version of this manuscript focused on the age effects; SS and MNR co-wrote the current version of the manuscript.

## Supplementary Materials

**Supplementary Table 1.**
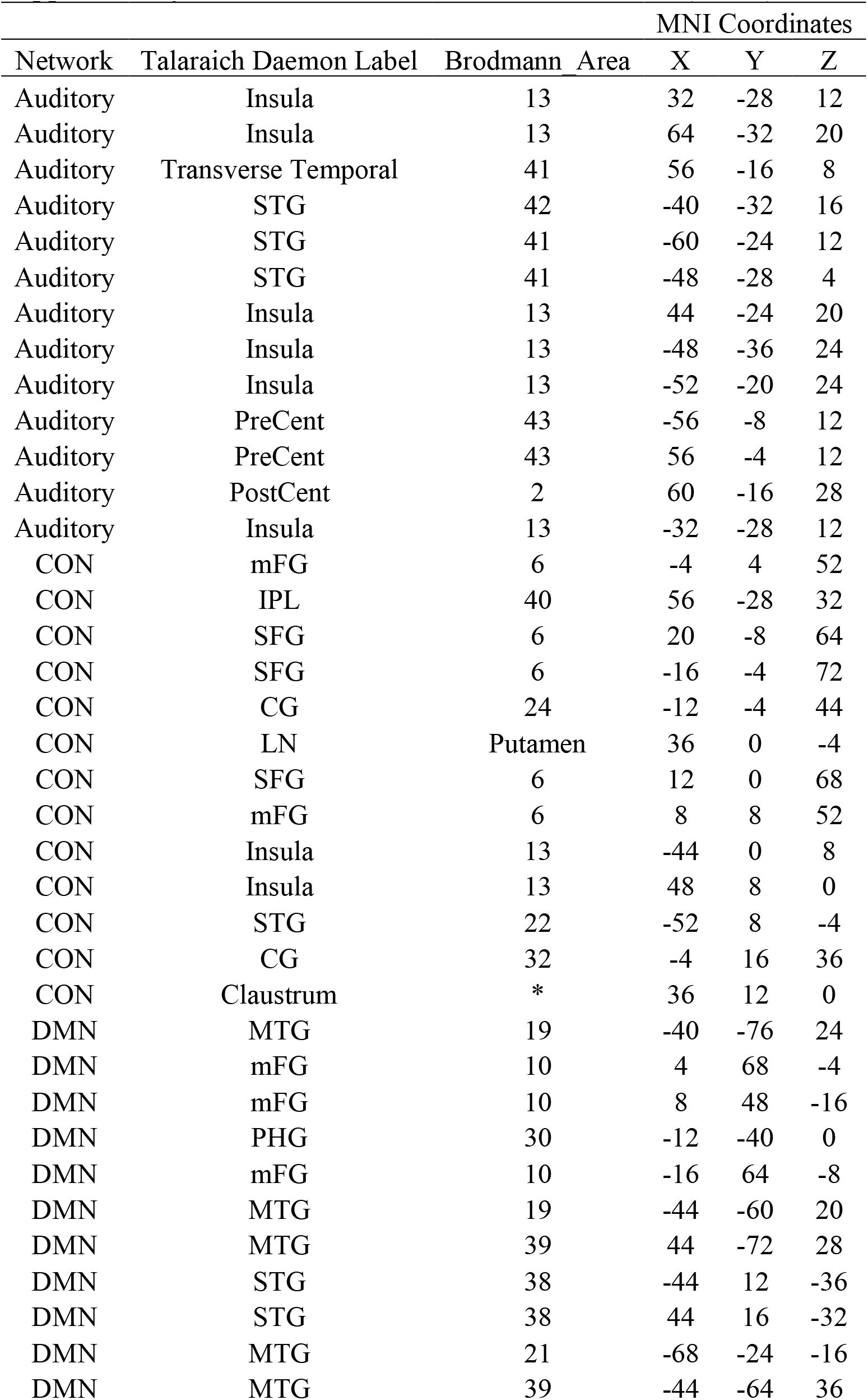
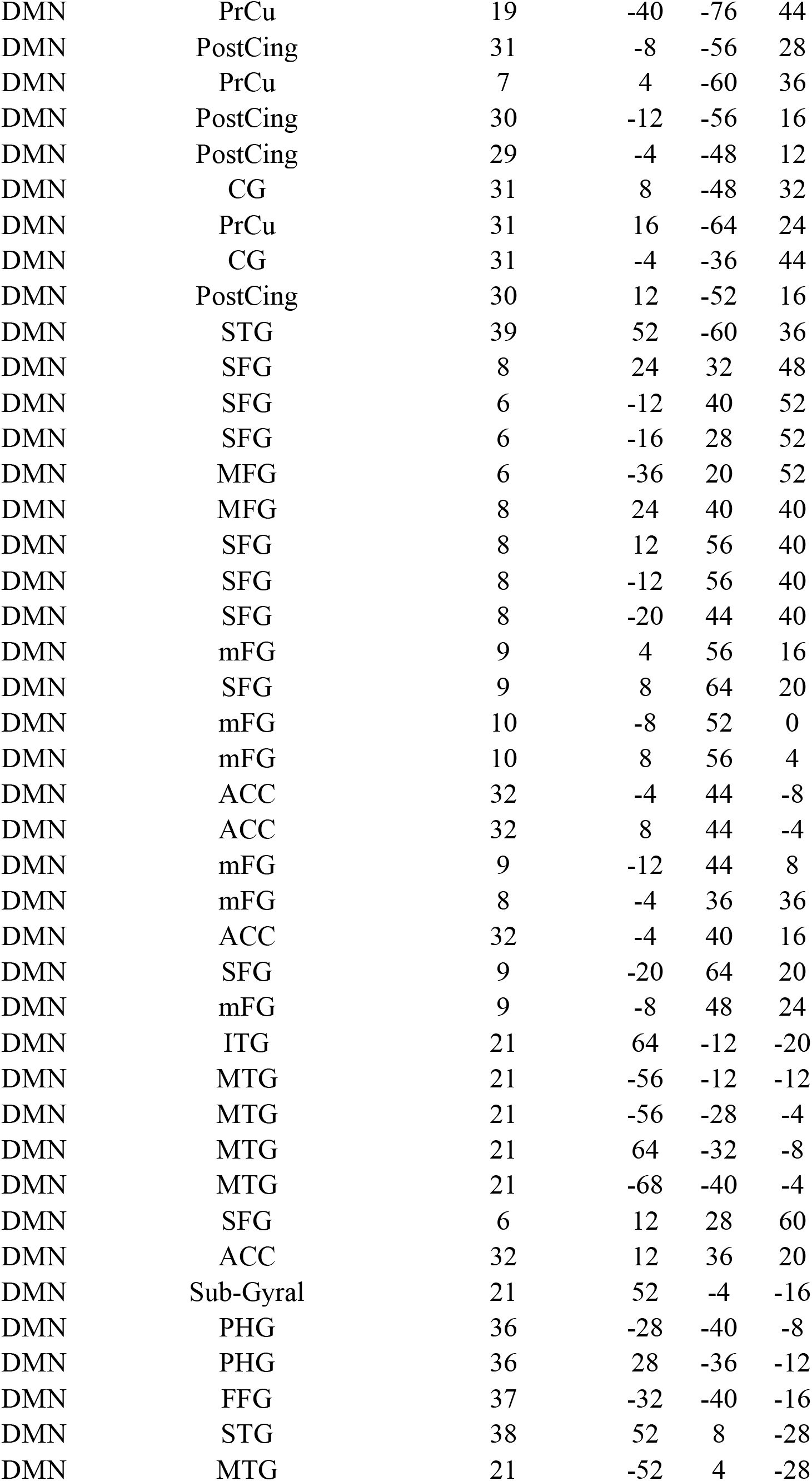
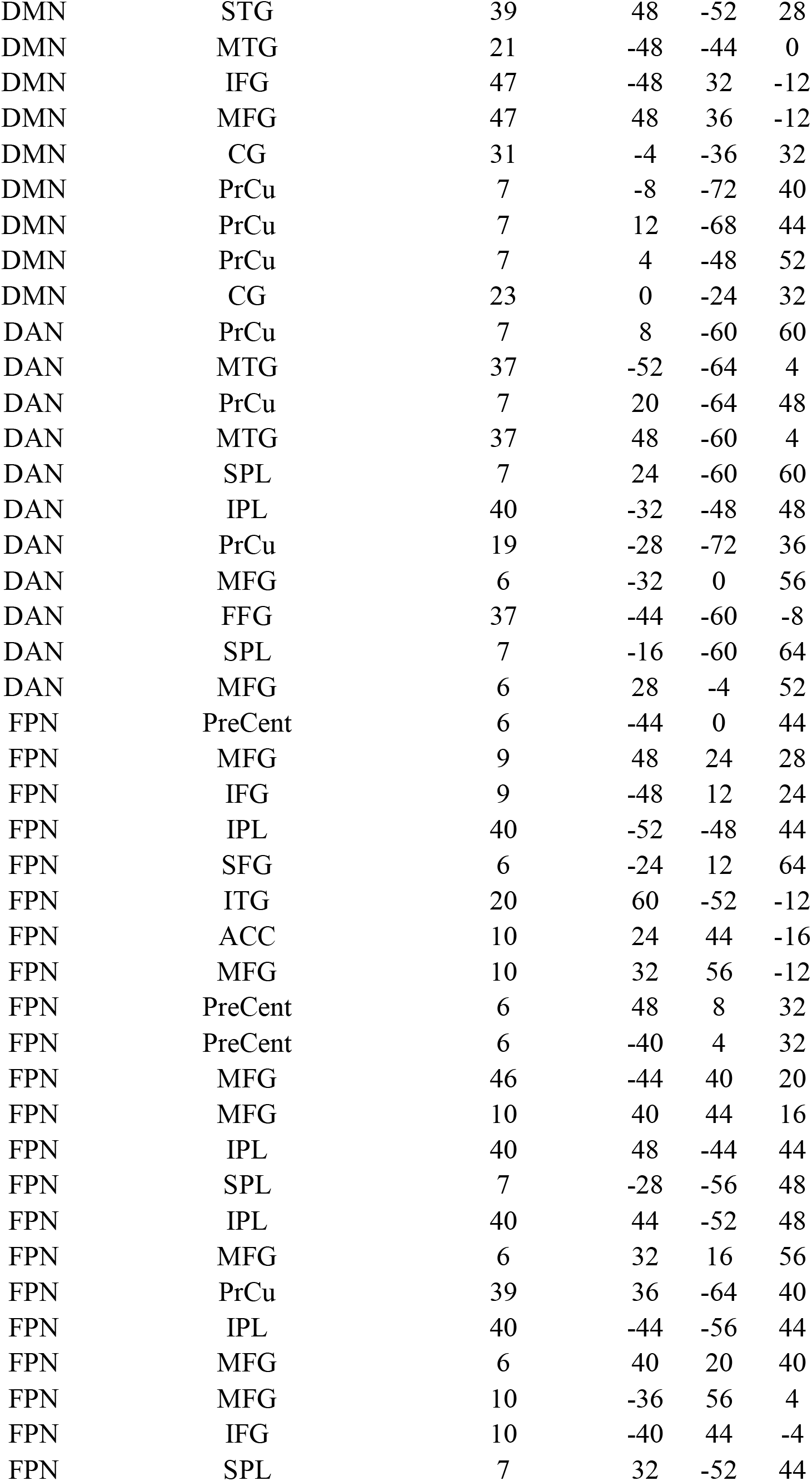
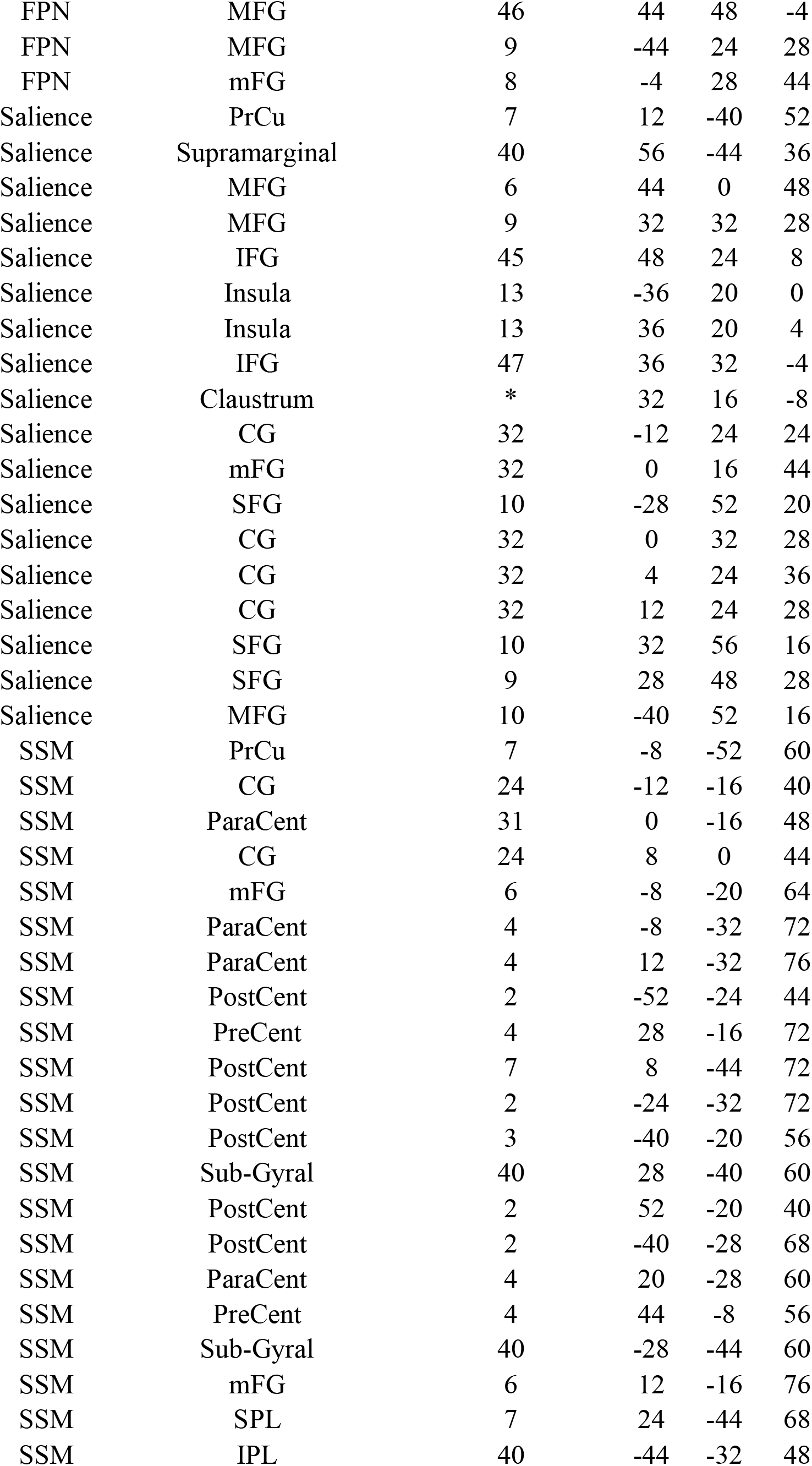
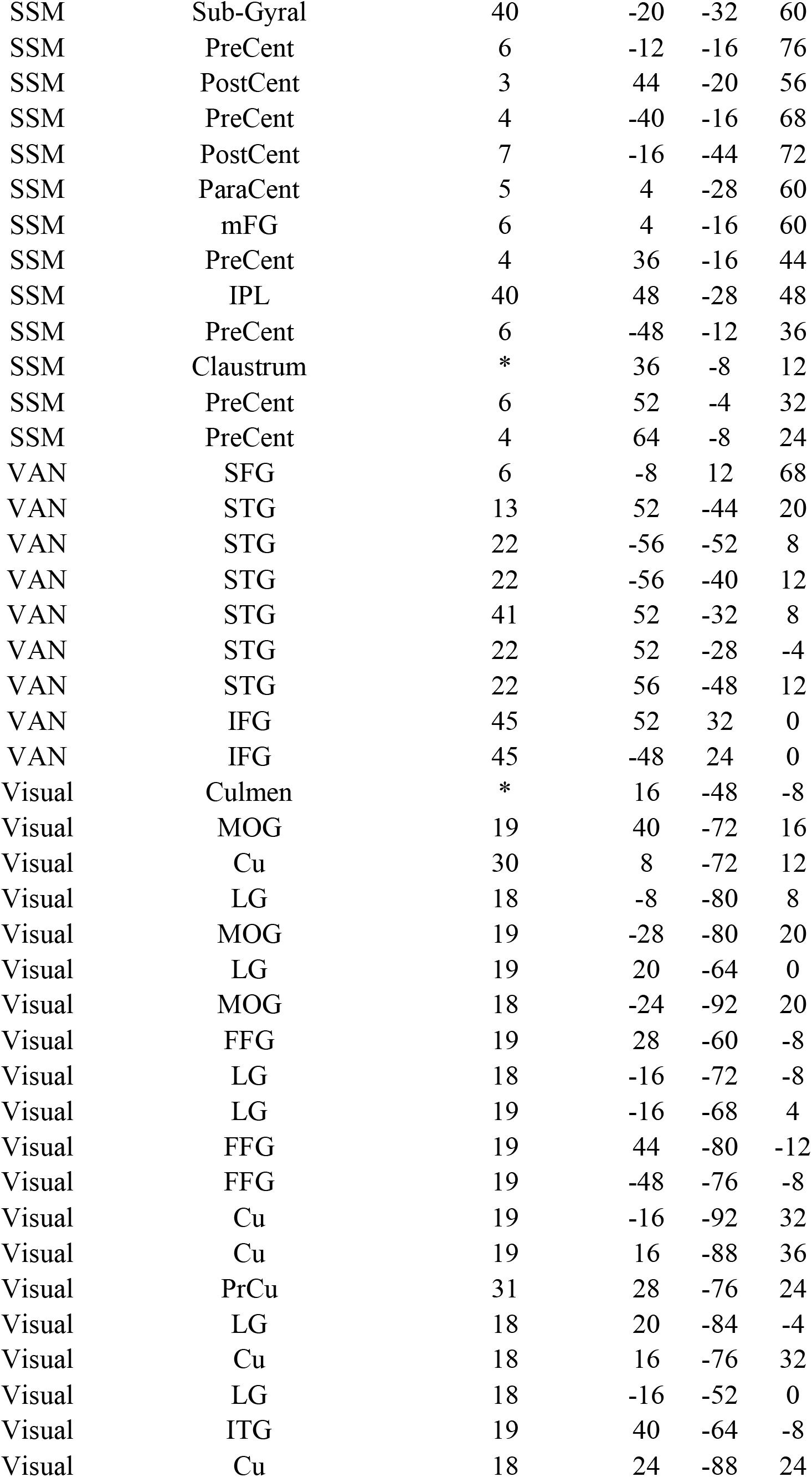
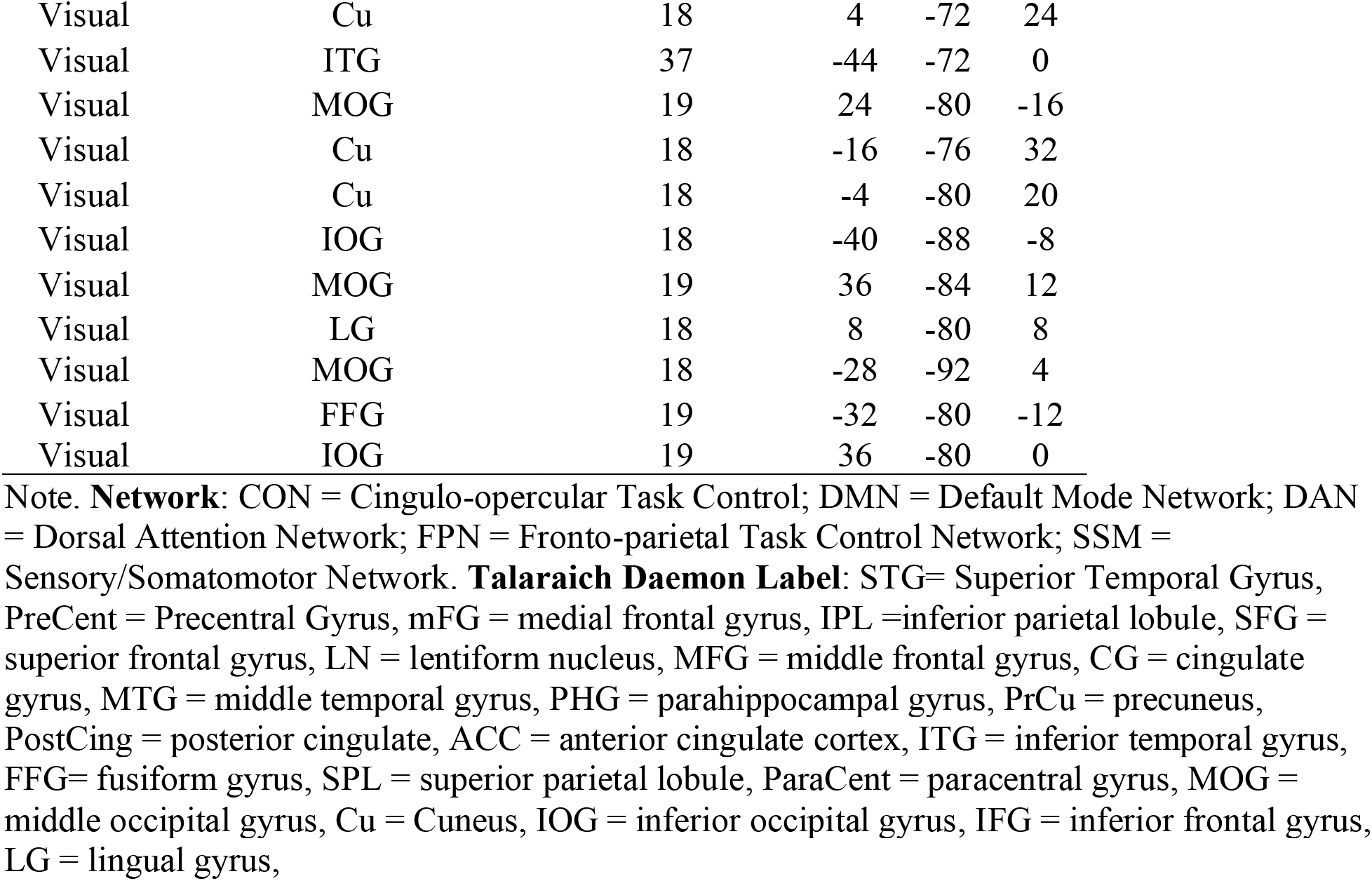
Power atlas ROI network nodes (n=216) used in the analysis.

### B-PLS2 Summary of Findings for LV3 and LV4

#### LV3: Sex similarities in accuracy-related, but differences in age-related, task connectivity

LV3 accounted for 11.45% of the cross-block covariance and identified significant positive correlation coefficients between several networks. The loading and density matrices (Suppl. Figure 1A and 1C, 1D) indicate that there was greatest density of positive correlations between the FPN and Salience networks. In addition, there were dense within-network positive correlations in the Salience and Visual networks. The loading matrix (Suppl. Figure 1A) and behavioral weights (Suppl. Figure 1B) indicate that in men, age was negatively correlated with increased connectivity in the aforementioned networks, and memory performance during both the SE and SH tasks was also negatively correlated with increased functional connectivity in these networks. In women, age was not significantly correlated with connectivity among these networks, and just like men memory performance for both SE and SH tasks was negatively correlated with increased functional connectivity in these networks. LV3 also identified negative correlations within DAN and between DAN, FPN and CON and Salience networks; and, between Visual-FPN, Salience and CON, and VAN-CON (Suppl. Figure 1D). These patterns of within- and between-network connectivity were correlated with advanced age in men, better memory performance both tasks in men and women.

**Supplementary Figure 1.**
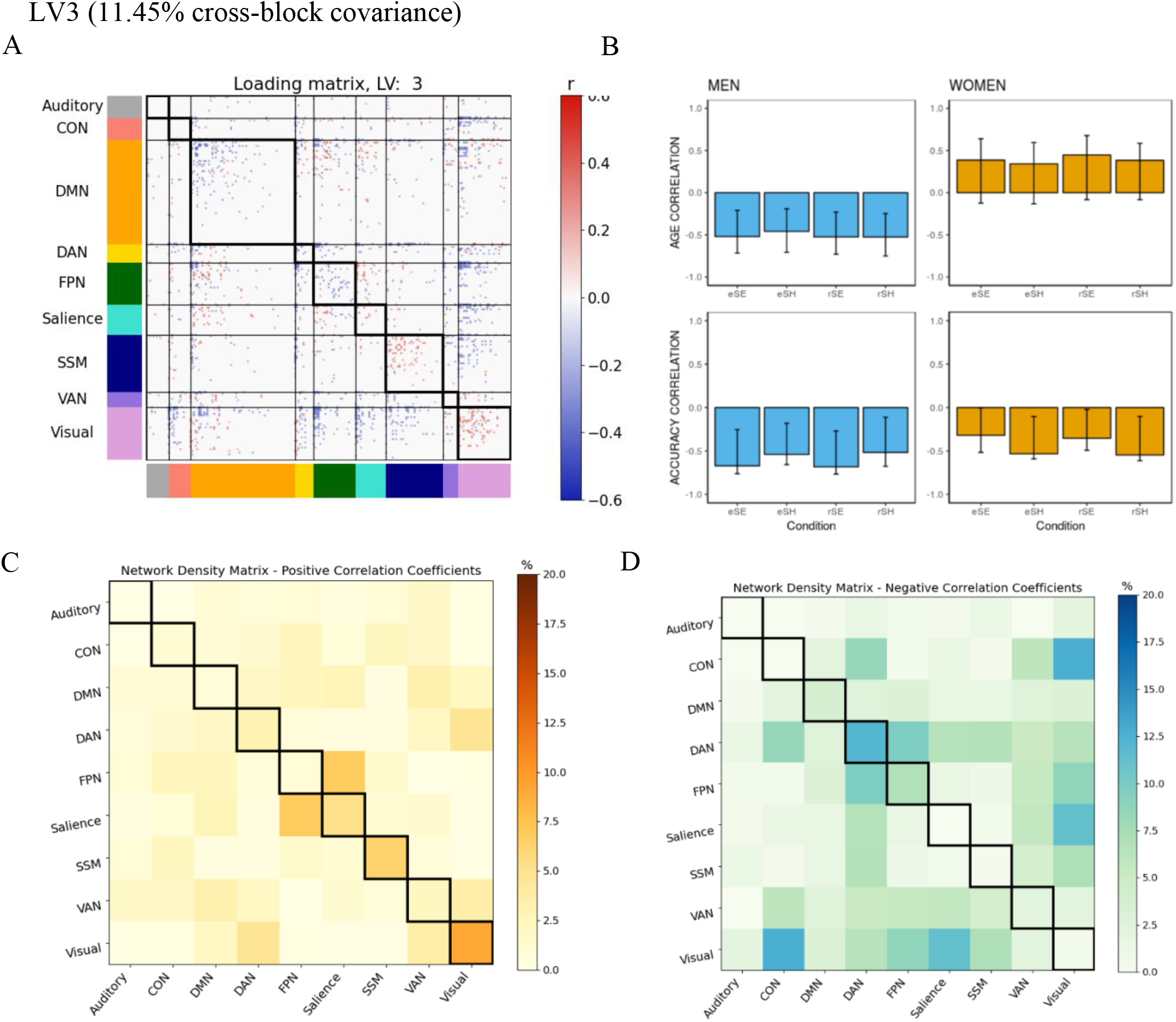
B-PLS2, LV3: Sex similarities in accuracy-related, but differences in age-related, task connectivity (Between-Sex Group B-PLS2 analysis) **(A)** Thresholded 95th percentile of correlations between participants’ task fMRI data and behavioral profile indicated in B. **(B)** Correlation between the behavioral vectors of age and accuracy with the task fMRI connectivity of participants (behavior correlation weights). Error bars represent bootstrapped standard deviations. **(C)** The density plot for the positive correlation coefficients (i.e., sum of the significant correlation coefficients after thresholding, divided by the total number of edges between any two networks). **(D)** The density matrix for the negative correlation coefficients. eSE = encoding spatial easy; eSH = encoding spatial hard; rSE = retrieval spatial easy; rSH = retrieval spatial hard; CON = cingulo-opercular network; DMN = default mode network; DAN = dorsal attention network; FPN = frontoparietal network; SSM = somatomotor network; VAN = ventral attention network.

#### LV4: Sex differences in accuracy, but no effects of age, in task connectivity

LV4 accounted for 5.23% of the cross-block covariance and showed only significant negative correlations. The loading and density matrices (Suppl. Figure 1E and 1G) show significant negative correlation connections between DAN-DMN and FPN, between SSM-CON, DMN and DAN and between Visual and FPN networks. Together with the brain-behavior plots (Suppl. Figure 1F), these networks show a strong positive correlation with memory performance on the task for the SE conditions in men. Conversely, women show a strong negative correlation between connectivity and performance on both SE and SH tasks in these same networks.

**Supplementary Figure 2.**
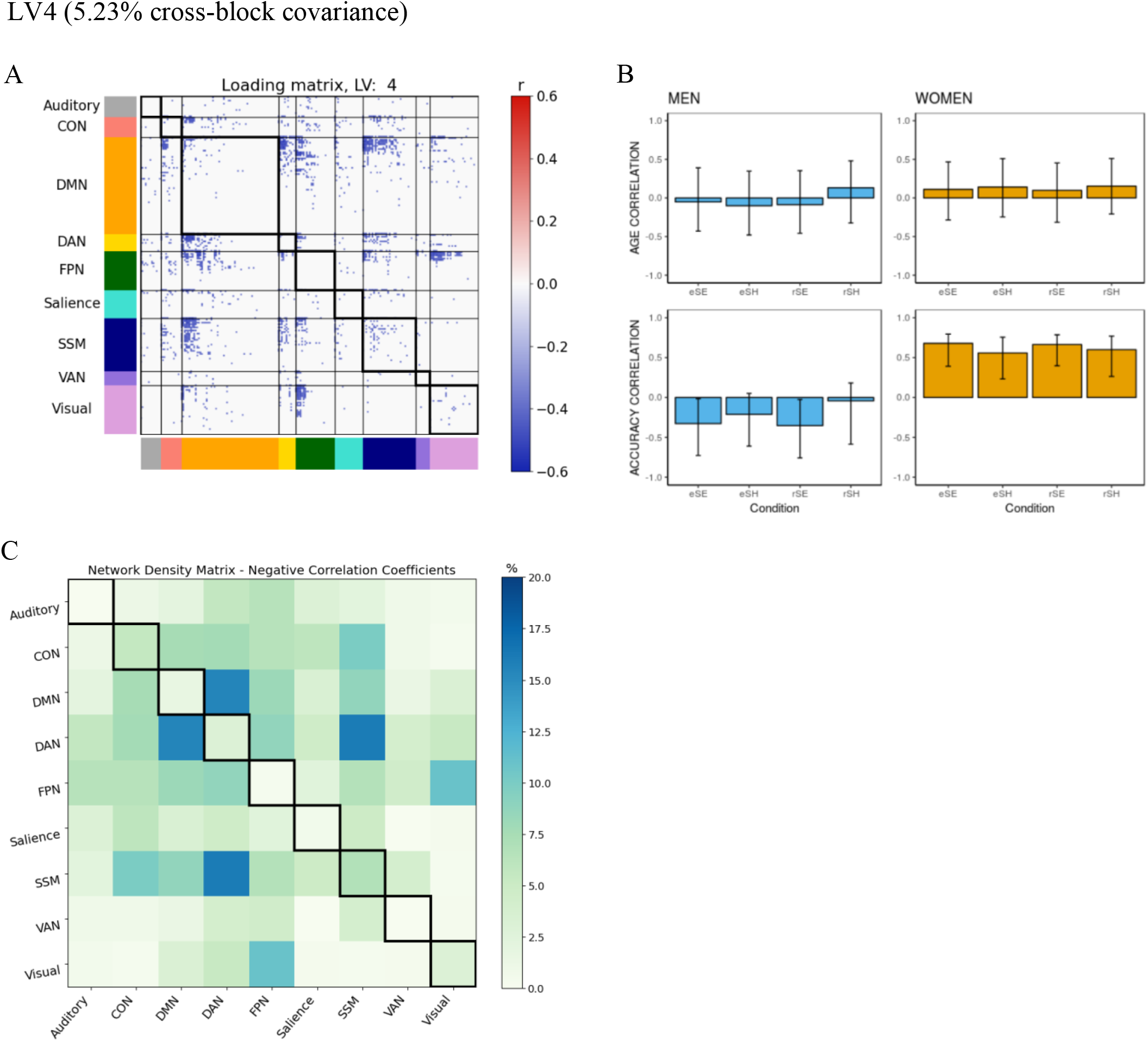
B-PLS2, LV4: Sex differences in accuracy, but no effects of age, in task connectivity (Between-Sex Group B-PLS2 analysis) **(A)** Thresholded 95th percentile of correlations between participants’ task fMRI data and behavioral profile indicated in B. **(B)** Correlation between the behavioral vectors of age and accuracy with the task fMRI connectivity of participants (behavior correlation weights). Error bars represent bootstrapped standard deviations. **(C)** The density plot for the negative correlation coefficients (i.e., sum of the significant correlation coefficients after thresholding, divided by the total number of edges between any two networks). The density matrix for the positive correlation coefficients is not presented because there were no significant edges. eSE = encoding spatial easy; eSH = encoding spatial hard; rSE = retrieval spatial easy; rSH = retrieval spatial hard; CON = cingulo-opercular network; DMN = default mode network; DAN = dorsal attention network; FPN = frontoparietal network; SSM = somatomotor network; VAN = ventral attention network.

**Supplementary Figure 3.**
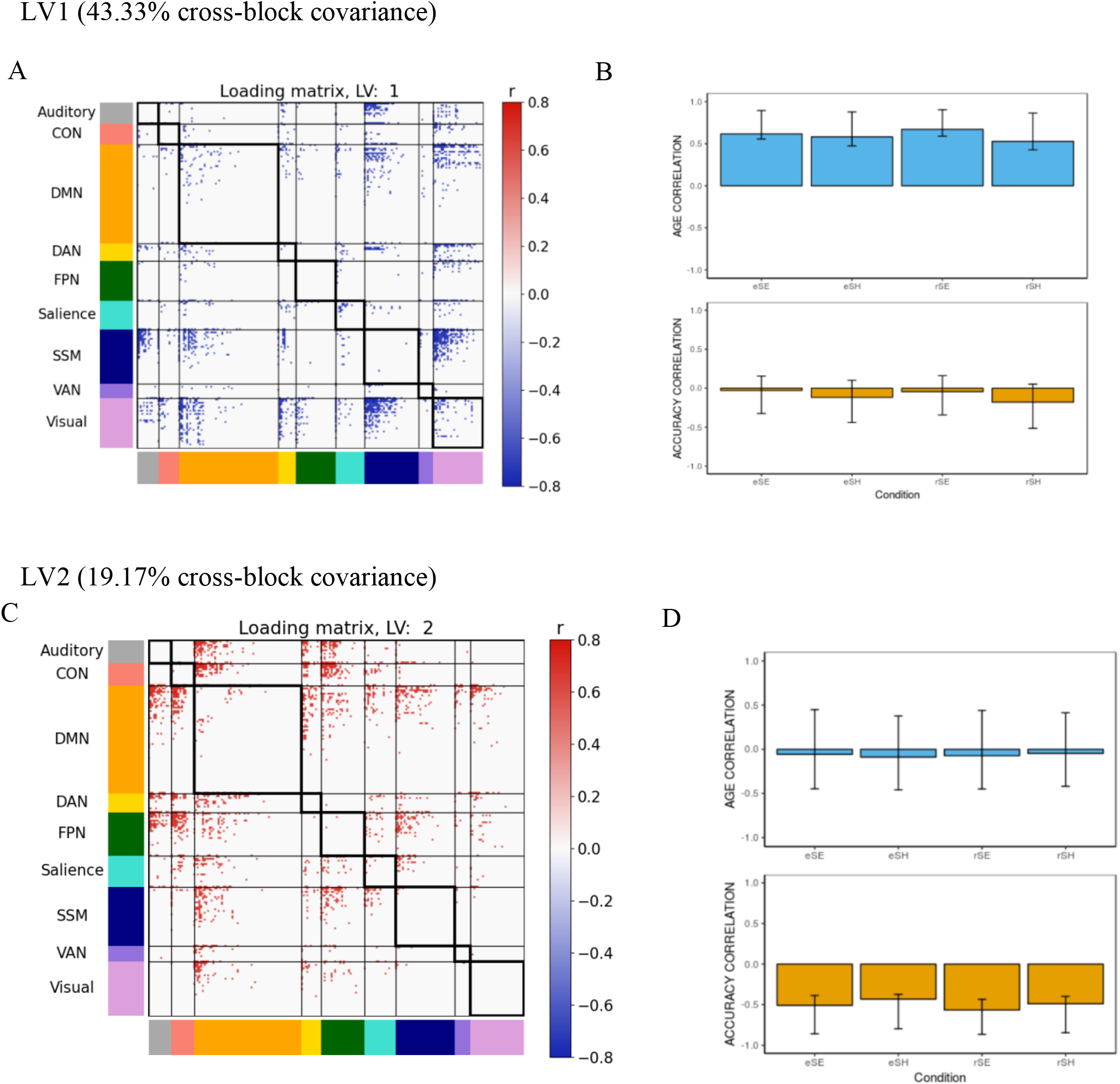
Matched - Sex cohort based on age, education, and intracranial volume (Full Group B-PLS1 Analysis) **(A, C)** thresholded 95th percentile of correlations between participants’ task fMRI data and behavioral profile for LV1 and LV2, respectively. (B, D) Behavioral profile of correlation between the behavioral vectors of age and accuracy with the task fMRI connectivity of participants (behavior correlation weights) for LV1 and LV2, respectively. Error bars represent bootstrapped standard deviations. eSE = encoding spatial easy; eSH = encoding spatial hard; rSE = retrieval spatial easy; rSH = retrieval spatial hard; CON = cingulo-opercular network; DMN = default mode network; DAN = dorsal attention network; FPN = frontoparietal network; SSM = somatomotor network; VAN = ventral attention network.

**Supplementary Figure 4.**
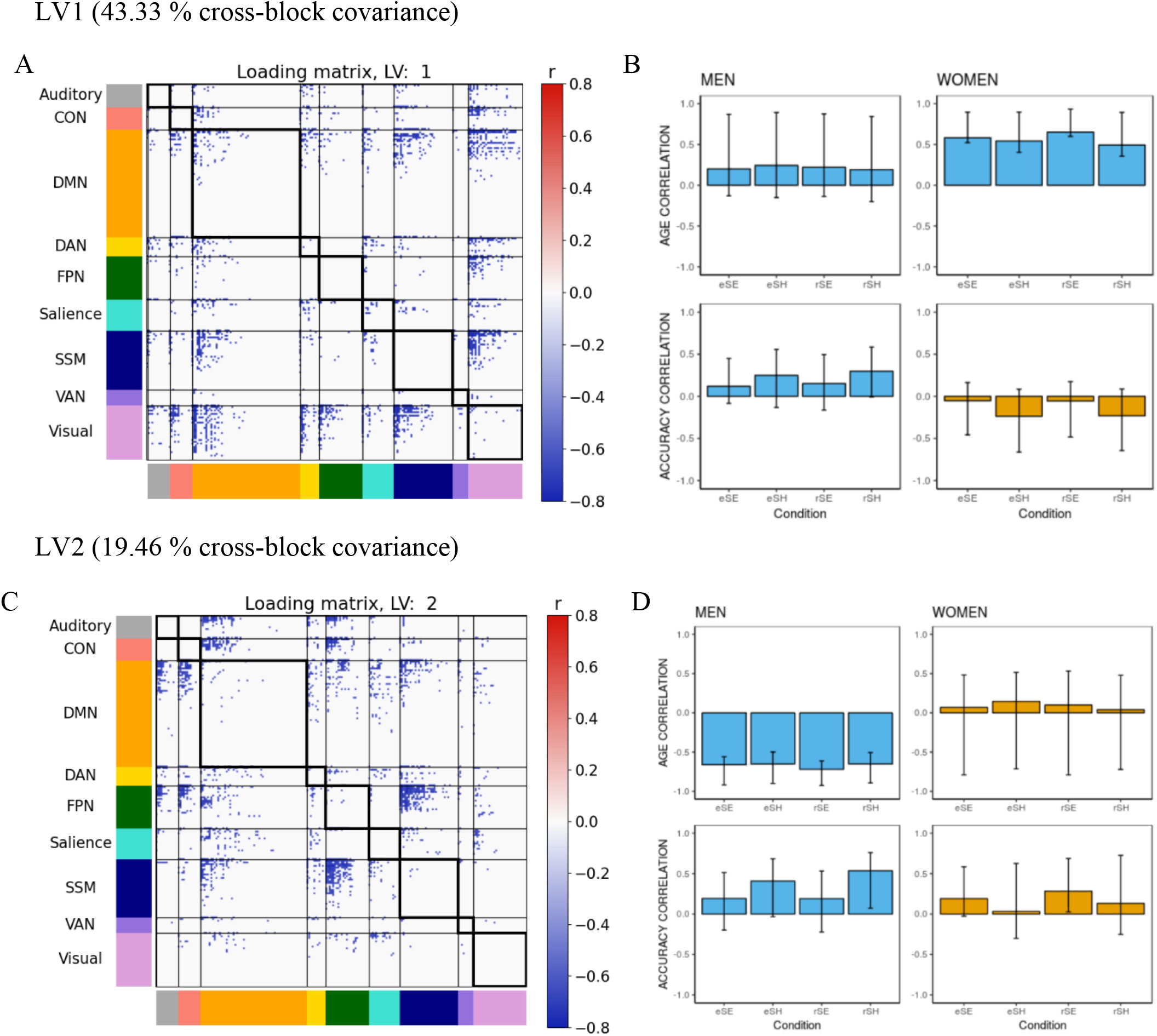
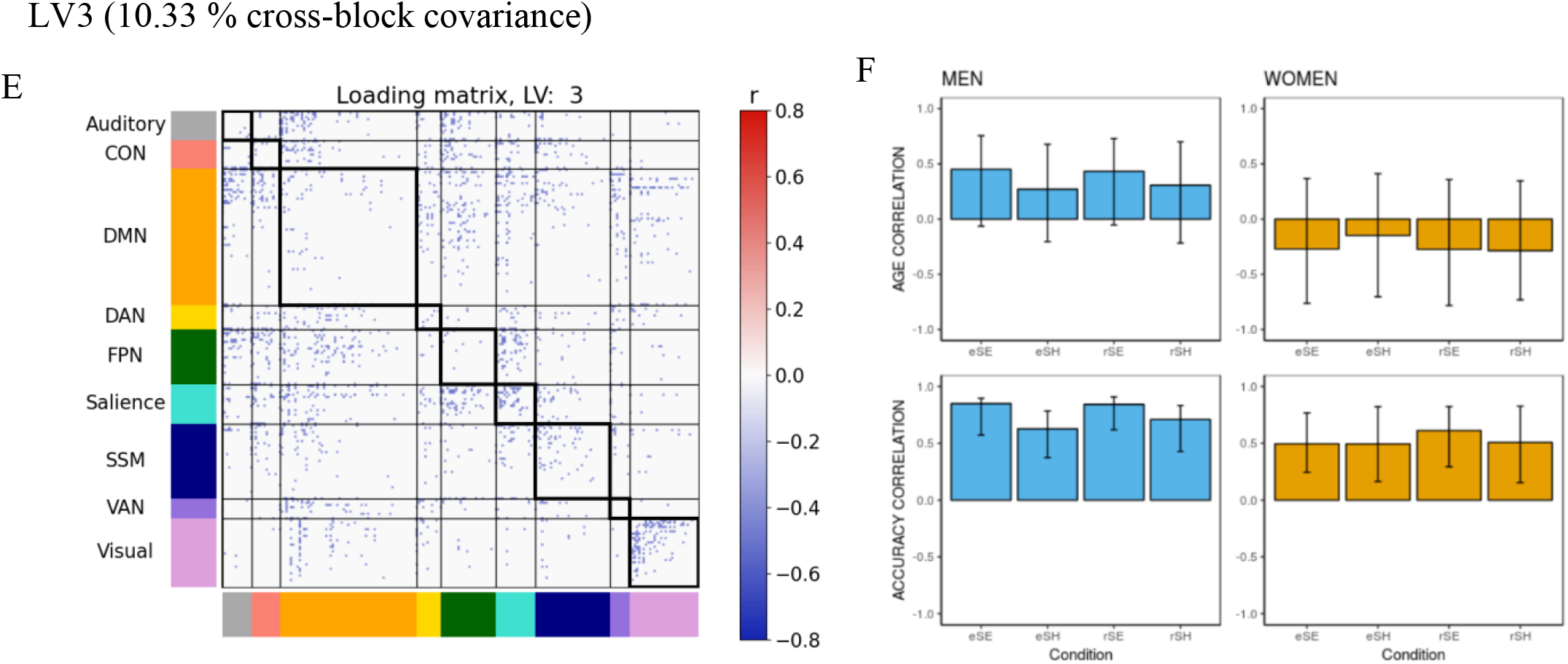
Matched-Sex cohort based on Age, Education, and Intracranial Volume (Between-Sex Group B-PLS2 Analysis) The BPLS analysis with a subcohort of participants matched by intracranial volume, age, and education (24 men, 24 women) within group (N=48) and between group (M=24, F=24) determined findings similar to the original BPLS analyses described in the manuscript. **(A, C, E)** thresholded 95th percentile of correlations between participants’ task fMRI data and behavioral profile for LV1, LV2, and LV3 respectively. **(B, D, F)** Behavioral profile of correlation between the behavioral vectors of age and accuracy with the task fMRI connectivity of participants (behavior correlation weights) for LV1, LV2, and LV3 respectively. Error bars represent bootstrapped standard deviations. eSE = encoding spatial easy; eSH = encoding spatial hard; rSE = retrieval spatial easy; rSH = retrieval spatial hard; CON = cingulo-opercular network; DMN = default mode network; DAN = dorsal attention network; FPN = frontoparietal network; SSM = somatomotor network; VAN = ventral attention network.

**Supplementary Figure 5.**
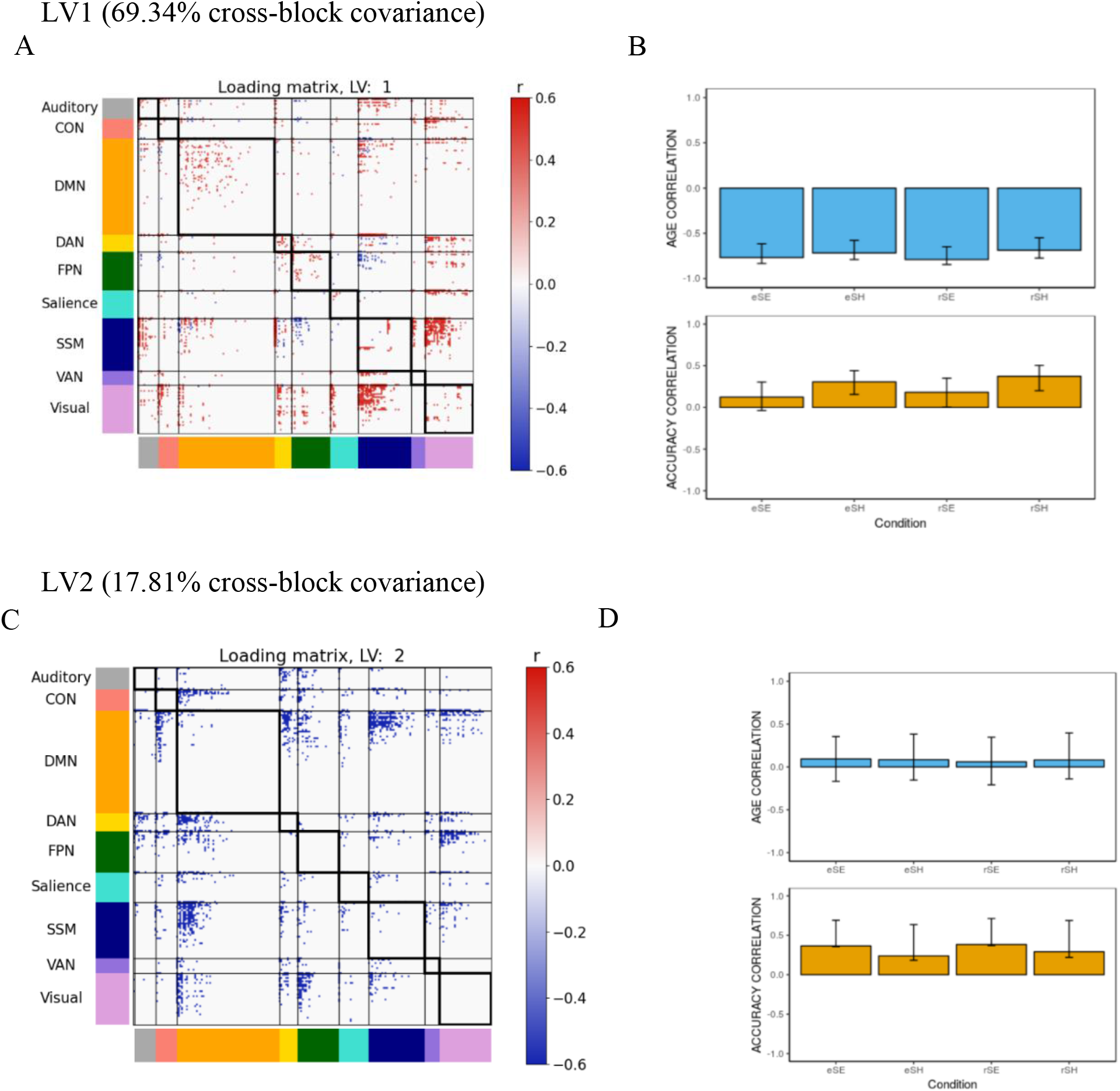
B-PLS1 without regressing mean task-related activity (Full Group B-PLS1 analysis) **(A, C)** thresholded 95th percentile of correlations between participants’ task fMRI data and behavioral profile for LV1 and LV2, respectively. **(B, D)** Behavioral profile of correlation between the behavioral vectors of age and accuracy with the task fMRI connectivity of participants (behavior correlation weights) for LV1 and LV2, respectively. Error bars represent bootstrapped standard deviations. eSE = encoding spatial easy; eSH = encoding spatial hard; rSE = retrieval spatial easy; rSH = retrieval spatial hard; CON = cingulo-opercular network; DMN = default mode network; DAN = dorsal attention network; FPN = frontoparietal network; SSM = somatomotor network; VAN = ventral attention network.

**Supplementary Figure 6.**
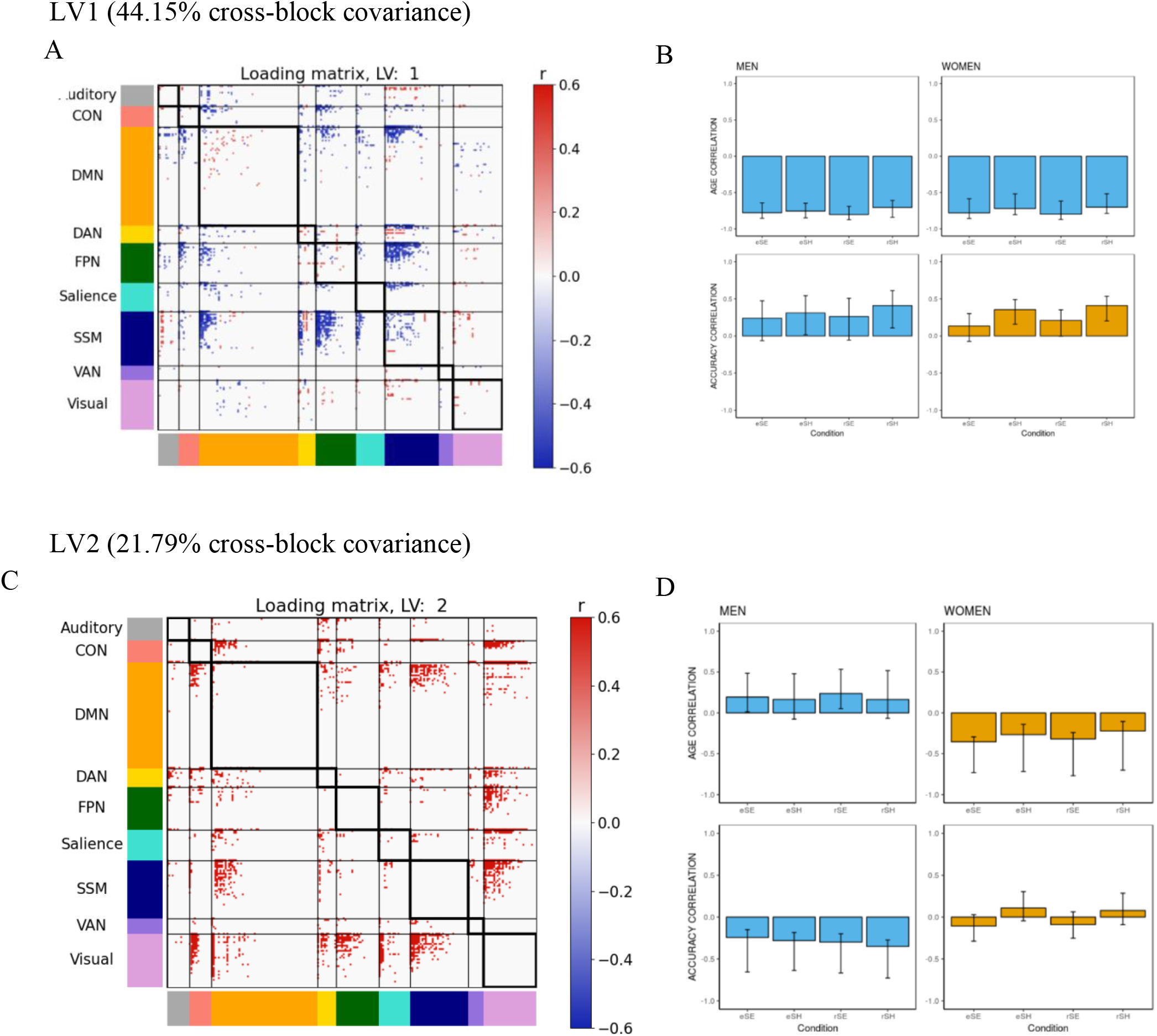
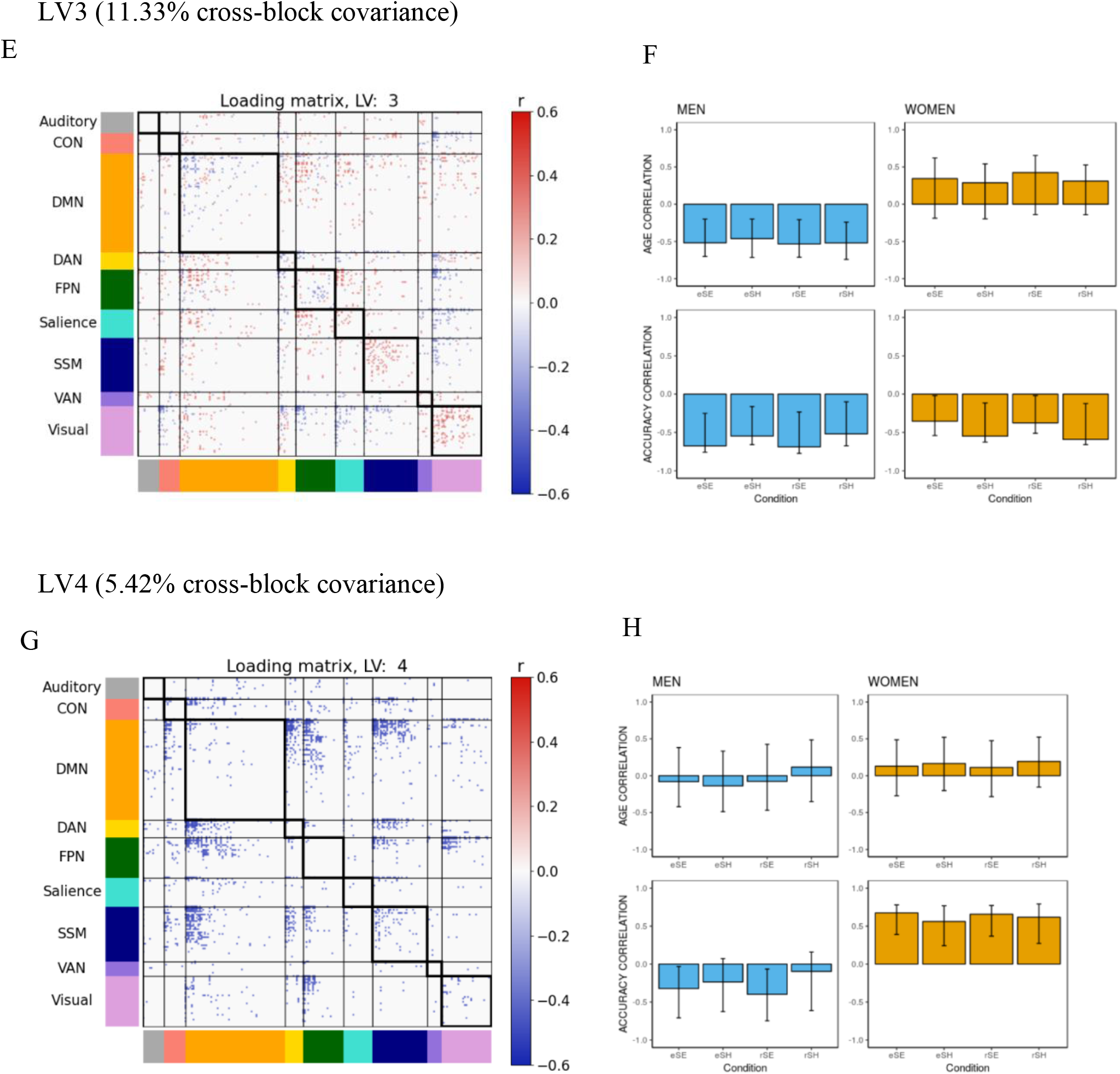
B-PLS2 without regressing mean task-related activity (Between-Sex Group B-PLS2 analysis) Note. The B-PLS analysis using connectivity matrices that did not regress mean-task-related activity generated 2 significant LVs within group and 4 significant LVs between group, similar to the primary BPLS (Analyses 1 & 2) described in the manuscript. **(A, C, E, G)** thresholded 95th percentile of correlations between participants’ task fMRI data and behavioral profile for LVs 1-4, respectively. **(B, D, F, H)** Behavioral profile of correlation between the behavioral vectors of age and accuracy with the task fMRI connectivity of participants (behavior correlation weights) for LVs 1-4 respectively. Error bars represent bootstrapped standard deviations. eSE = encoding spatial easy; eSH = encoding spatial hard; rSE = retrieval spatial easy; rSH = retrieval spatial hard; CON = cingulo-opercular network; DMN = default mode network; DAN = dorsal attention network; FPN = frontoparietal network; SSM = somatomotor network; VAN = ventral attention network.

